# Erythroid overproduction of erythroferrone causes iron overload and developmental abnormalities in mice

**DOI:** 10.1101/2021.01.09.426054

**Authors:** Richard Coffey, Grace Jung, Joseph D. Olivera, Gabriel Karin, Renata C. Pereira, Elizabeta Nemeth, Tomas Ganz

**Author notes:** Corresponding Author: Tomas Ganz, PhD, MD.

## Abstract

The hormone erythroferrone (ERFE) is produced by erythroid cells in response to hemorrhage, hypoxia or other erythropoietic stimuli, and suppresses the hepatic production of the iron-regulatory hormone hepcidin, thereby mobilizing iron for erythropoiesis. Suppression of hepcidin by ERFE is thought to be mediated by interference with paracrine BMP signaling that regulates hepcidin transcription in hepatocytes. In anemias with ineffective erythropoiesis, ERFE is pathologically overproduced but its contribution to the clinical manifestations of these anemias is not well understood. We generated three lines of transgenic mice with graded erythroid overexpression of ERFE and showed that they developed dose-dependent iron overload, impaired hepatic BMP signaling and relative hepcidin deficiency. These findings add to the evidence that ERFE is a mediator of iron overload in conditions where ERFE is overproduced, including anemias with ineffective erythropoiesis. At the highest levels of ERFE overexpression the mice manifested decreased perinatal survival, impaired growth, small hypofunctional kidneys, decreased gonadal fat depots and neurobehavioral abnormalities, all consistent with impaired organ-specific BMP signaling during development. Neutralizing excessive ERFE in congenital anemias with ineffective erythropoiesis may not only prevent iron overload but may have additional benefits for growth and development.

**Key Points:**

1. Chronic erythroid overproduction of erythroferrone dose-dependently suppresses hepcidin, causing iron overload even in the absence of anemia
2. High level overexpression of erythroferrone can cause delayed growth, impaired kidney function and other developmental abnormalities consistent with altered BMP signaling

## Introduction

Systemic iron homeostasis is regulated by the hepatic hormone hepcidin, which inhibits intestinal iron absorption and the mobilization of stored iron^1^. Erythropoietic activity, dependent on adequate iron supply, strongly influences iron homeostasis. During the physiological response to anemia, within hours after erythropoietin (EPO) induction, hepcidin is suppressed^2^, increasing iron flows into blood plasma and iron delivery to the marrow to support the enhanced production of erythrocytes. In anemias with ineffective erythropoiesis, in which EPO levels are increased and the erythropoietic compartment is disproportionately expanded, hepcidin is pathologically suppressed^3,4^ resulting in excessive iron absorption and eventual iron overload, even in untransfused patients. The search for factors that suppress hepcidin in response to increased erythropoiesis^5^ led to the discovery of erythroferrone (ERFE), a hormone produced by erythroblasts that acts on the liver to suppress hepcidin expression by hepatocytes^6^.

Recent studies provided evidence that ERFE inhibits hepcidin expression by antagonizing select members of the bone morphogenetic protein (BMP) family^7,8^. BMP2/6 signaling in hepatocytes regulates hepcidin transcription in response to plasma iron concentrations and hepatic iron stores, and the pathological loss of BMP signaling causes hepcidin deficiency and systemic iron overload^9,10^. ERFE lowers hepcidin transcription by sequestering the BMP2/6 heterodimer secreted by the sinusoidal endothelial cells ^8^, but ERFE is also reported to bind other BMPs^7,11^. Considering the pleiotropic effects of BMPs, the endocrine effects of pathologically-increased ERFE need not be limited only to altered hepcidin regulation in the liver.

Although substantial evidence supports the role of ERFE as a stress hormone that acutely suppresses hepatic hepcidin synthesis in response to erythropoietic stimuli, much less is known about the chronic effects of ERFE *in vivo*, which have been analyzed only in the context of erythropoietic disorders such as β-thalassemia^12^. While patients with β-thalassemia commonly suffer from problems that could be linked to disruptions in BMP signaling, including iron overload^13^, renal impairment^14^, and skeletal problems such as impaired growth^15^ or bone mineralization^16^, it is unclear to which extent the observed pathologies are attributable to elevated ERFE as opposed to other factors such as tissue and organ hypoxia, the effects of hemolytic products or the toxicity of treatment. Animal models of increased ERFE production depend on the stimuli of anemia^17,18^ or EPO administration ^19^ but serum ERFE levels in these models could be orders of magnitude lower than in human patients with β-thalassemia^12,20^. Additionally, high EPO levels exert multiple systemic effects^21^ beyond increasing *Erfe* expression and these complicate direct attribution of observed phenotypes to the action of ERFE alone.

The aim of the current study was to determine the effect of chronically elevated *Erfe* expression on systemic iron homeostasis and BMP signaling, without the confounding effects of chronic anemia or EPO treatment, and at a wide range of circulating ERFE levels to include those present in humans with β-thalassemia and other anemias with ineffective erythropoiesis. To this end we generated multiple lines of novel transgenic mice overexpressing graded levels of erythroferrone in erythroid cells. We find that chronically elevated *Erfe* expression results in dose-dependent hepcidin suppression, dysregulation of iron homeostasis and tissue iron accumulation. Additionally, at high levels, chronic *Erfe* overexpression affects multiple organ systems influencing pup survival, somatic growth, fat depots, kidney function, and behavior.

## Materials and Methods

### Transgenic mice

A DNA construct targeting gene overexpression to erythroid cells was generously contributed by Dr. Kenneth R. Peterson ^22^ and engineered to express N-FLAG-*mErfe* **(Figure 1A)**. Transgenic founder animals were generated at Cyagen labs by pronuclear injection of fertilized C57BL/6N zygotes. One transgenic founder gave rise to the high-expressing “H” line and the moderately expressing “M” line, and the second transgenic founder gave rise to the low-expressing “L” line. Individual transgenic lines were established and experimental mice generated by breeding heterozygous transgenic mice with C57BL/6J mice to yield transgenic mice and line-specific WT littermate controls. Each mouse was genotyped to ensure stability of transgene copy number.

**Figure 1:**
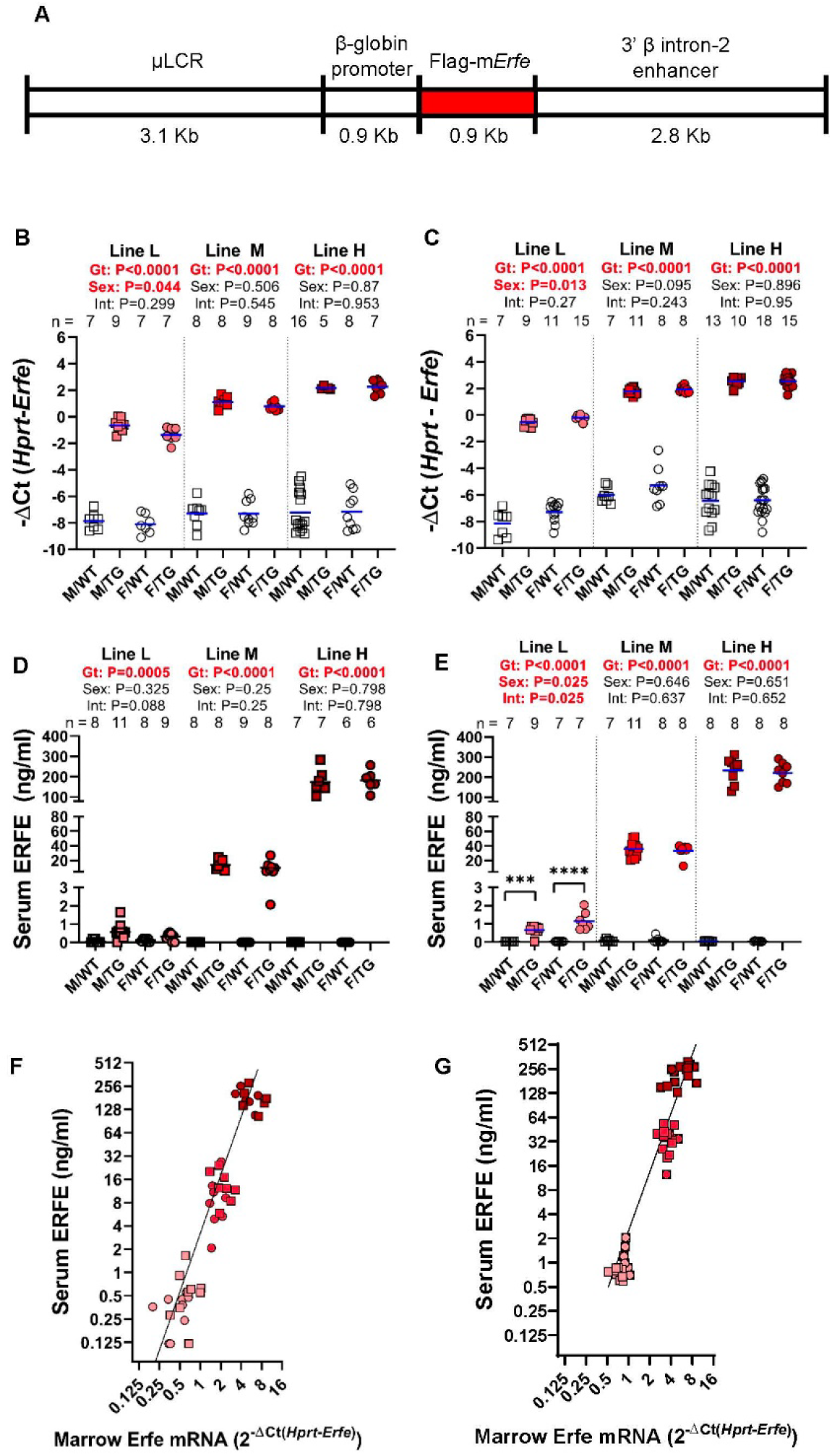
Generation of Hbb-*Erfe* mice and analysis of *Erfe* overexpression. **A)** Structure of the transgene construct. Bone marrow *Erfe* mRNA in **B**) 6 week-old and **C**) 16-week old mice. Serum ERFE levels of **D)** 6-week-old and **E)** 16-week-old mice. Relationship between serum ERFE protein levels and marrow *Erfe* mRNA in **F)** 6-week-old mice, regression line Y=3.18* X^2.5^ (r^2^ =0.51), and **G)** 16-week-old mice, regression line Y=0.83* X^2.4^ (r^2^ =0.57). In all panels, male (M, square) and female (F, circle); mean of each group is shown by blue line, TG =*Erfe*-overexpressing mice (line-L pink, line-M red, and line-H dark red symbols) and wild-type = WT littermate controls (white symbols). Within each individual transgenic line, groups were compared by two-way ANOVA to determine significant effects (P<0.05, bold red) of genotype (**Gt**) and sex (**Sex**) on data variation and to identify interactions (**Int**) between these variables. Cohort numbers for mRNA and serum analysis are shown.

Routine Materials and Methods are in Supplemental Data.

## Results

### Generation of Erfe-overexpressing mice

The three lines of *Erfe*-transgenic mice, referred to as high (line-H), medium (line-M), and low (line-L), differed in the magnitude of *Erfe* overexpression, but all had increased bone marrow *Erfe* expression relative to their line-specific wild-type (WT) littermates **(Figure 1B)**. Lines-H and -M had been derived from the highest-expressing founder. Genomic sequencing identified the haploid transgene insertion site on Chr. 4, from 11985998 to 12010497, overlapping into gene 1700123M08Rik noncoding RNA of unknown function. The H- and M-lines differed by transgene copy number (3 vs 1), resulting from a rare spontaneous deletion of 2 tandem copies of the transgene. Line-L was derived from another founder and was not further genomically characterized. Transgene expression was similar at 6- and 16-weeks **(Figure 1B, C)**.

At either age, serum ERFE levels in line-L mice were near the threshold of detection by ELISA, below levels previously measured in WT mice after EPO injection and similar to those of Hbb^Th3/+^ thalassemic mice ^18^ **(Figure 1D,E and Supplemental Figure 1)**. Compared with line-L, serum ERFE levels were ~20× higher in line-M and ~200× in line-H mice. Interestingly, moderate differences in mRNA bone marrow *Erfe* expression led to much larger differences in circulating ERFE levels between transgenic lines at either 6 or 16 weeks of age **(Figure 1F,G)**.

### Erfe overexpression causes iron overload and elevated hemoglobin levels

*Erfe* overexpression dose-dependently increased liver non-heme iron levels. At 6 weeks, mice from lines L, M and H respectively loaded ~2, ~3.6 and ~4.2 times more liver non-heme iron than their WT littermates **(Figure 2A)**. The effect of elevated *Erfe* expression on liver iron accumulation persisted in 16-week-old transgenic mice **(Figure 2B)**. Females had higher liver iron levels than males, consistent with the previously described sex differences in iron loading of the C57BL/6 strain ^23^. Serum iron was also elevated in mice from line-M and H, but not line-L, at 6 weeks **(Figure 2C)**, and increased in 16-week-old transgenic mice in all lines relative to WT littermates **(Figure 2D)**. We characterized in detail the pattern of iron loading in line-H mice at 16 weeks. Histochemical analysis using enhanced Perls stain showed that hepatic iron accumulation was periportal (**supplemental Figure 2A**). Liver *Tfrc* mRNA expression was decreased, consistent with increased hepatic iron loading, with no change in the mRNA expression of the hepcidin regulators *Tmprss6* or *Tfr2* (**supplemental Figure 2B-D**). Quantitation of non-hepatic tissue iron levels demonstrated significantly increased non-heme iron in the pancreas and heart of line-H mice, and a trend toward increased iron in the spleen (**supplemental Figure 2E-G**). Moreover, serum ferritin and transferrin saturation were increased compared with WT controls (**supplemental Figure 2H,I**).

**Figure 2:**
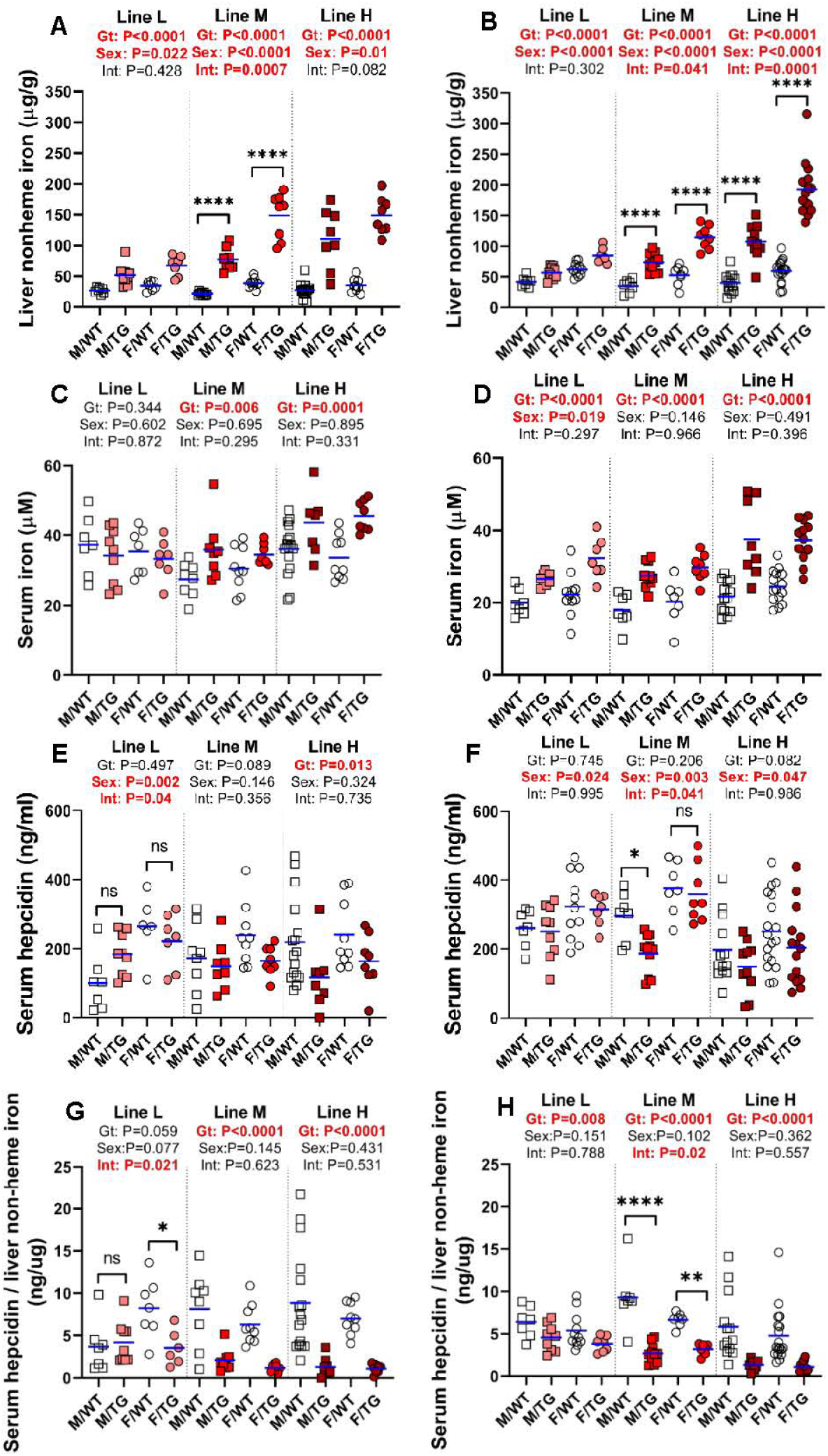
*Erfe* overexpression causes dose-dependent iron accumulation and inadequate hepcidin expression. Liver non-heme iron (**A, B**), serum iron (**C, D**), serum hepcidin (**E, F**), and serum hepcidin relative to liver iron (**G**, **H**) levels at 6 (**A, C, E, G**) and 16 (**B, D, F, H**) weeks of age in male (M) and female (F) *Erfe*-overexpressing (TG) mice and wild-type (WT) littermate controls from line-L (white/pink), line-M white/red), and line-H (white/dark red). The mean of each group is indicated by blue line. For each mouse line, groups were compared by two-way ANOVA to determine effects of genotype and sex on data variation (significant differences denoted in bold red) and to identify interactions between these variables. In the event of significant interaction between genotype and sex, individual groups were compared by Šidak’s multiple comparisons test (ns=P≥0.05, *=P<0.05, **=P<0.01, ***=P<0.001, ****=P<0.0001).

Serum hepcidin concentrations in *Erfe-*transgenic mice are expected to be determined by the balance between the suppressive effect of ERFE on hepcidin synthesis and the stimulatory effect of increased hepatic and plasma iron concentrations ^24,25^. In line-L mice at 6 or 16 weeks, serum hepcidin concentrations were similar to those of WT mice **(Figure 2E, F)**, consistent with these transgenic mice reaching iron balance by 6 weeks. In line M, the mean serum hepcidin concentrations were lower than their WT littermates but this reached statistical significance only in males at 16 weeks. In line-H, hepcidin was significantly suppressed in both sexes at 6 weeks and trended so at 16 weeks. Importantly, <u>the ratio</u> of serum hepcidin concentrations to liver non-heme iron content was lower in transgenic than WT mice at either 6 or 16 weeks, indicating inappropriately low hepcidin production relative to the severity of iron loading, with the exception of line-L males at 6 weeks **(Figure 2G, H)**.

At 6 and 16 weeks, line-H mice had elevated hemoglobin levels **(Figure 3A, B)** but not RBC counts **(Figure 3C, D)**, and their erythrocytes contained more hemoglobin **(Figure 3E, F)**. A similar effect was seen in line-M mice at 16 weeks but not 6 weeks. Line-L erythrocyte parameters were not different from WT.

**Figure 3:**
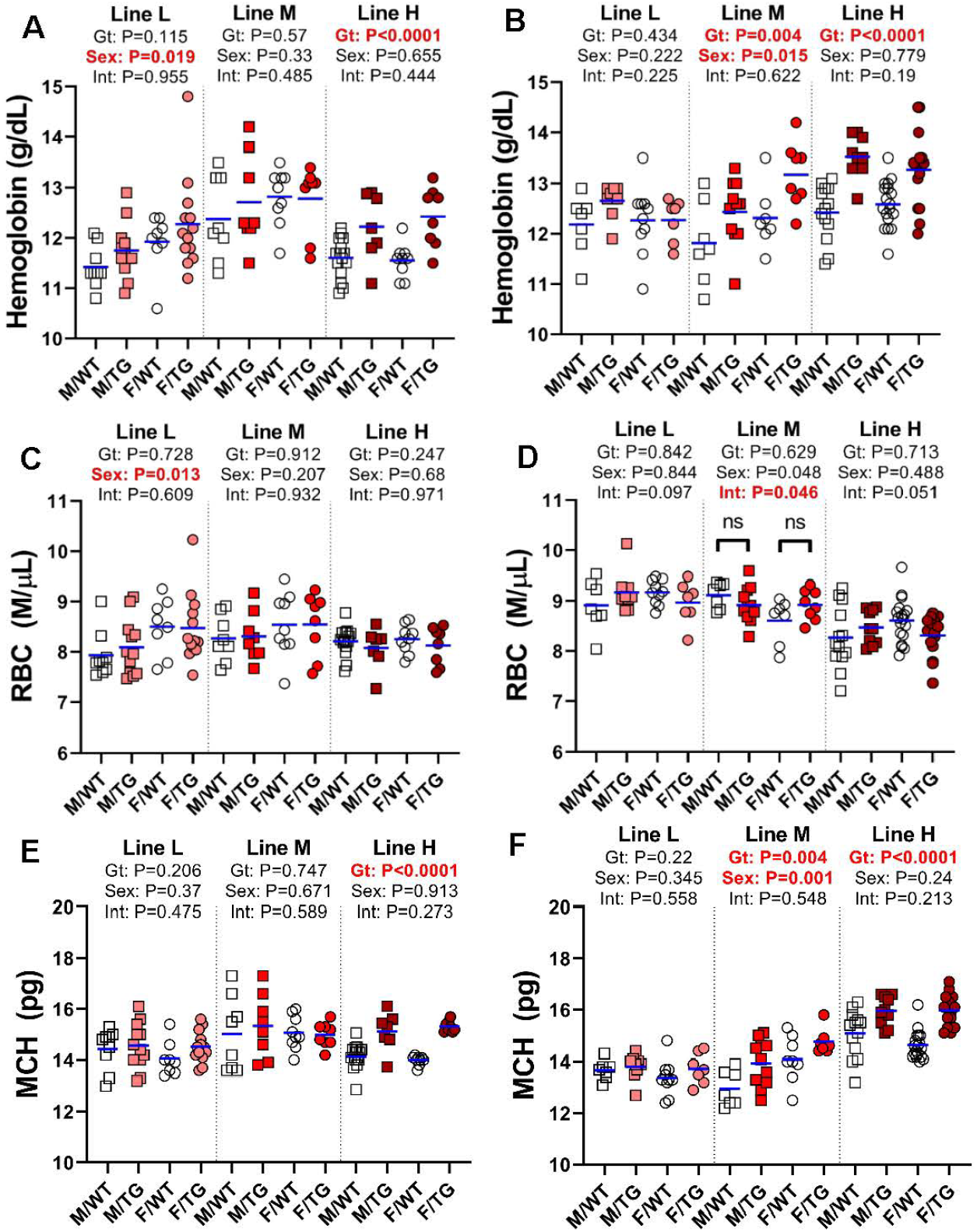
ERFE enhances hemoglobin synthesis in a dose-dependent manner. Hemoglobin levels (**A**, **B**), red blood cell (RBC) counts (**C, D**), and mean corpuscular hemoglobin (MCH) levels (**E**, **F)** at 6 (**A, C, E**) and 16 (**B, D, F**) weeks of age in male (M, square) and female (F, circle) wild-type (WT, white symbols) and *Erfe*-overexpressing (TG, colored symbols) mice from lines L (white/pink), M (white/red), and H (white/dark red). Group means are indicated by blue lines and groups within each individual line and age group were compared by two-way ANOVA to determine significant effects of genotype and sex on data variation and to identify interactions between these variables (P<0.05 denoted in bold red). In the event of significant interaction between genotype and sex, individual groups were compared by Šidak’s multiple comparisons test (NS=P>0.05, *=P<0.05, **=P<0.01, ***=P<0.001, ****=P<0.0001).

### Tissue-selective effect of Erfe overexpression on BMP signaling

In the liver, we detected a suppressive effect of higher ERFE concentrations on the hepatic expression of *Hamp* and *Smad7* but not *Id1* **(Figure 4A-C)**. There was no effect of high-level *Erfe* overexpression on the hepatic mRNA levels of *Bmp2* and *Bmp6* **(Figure 4D-E)**.

**Figure 4:**
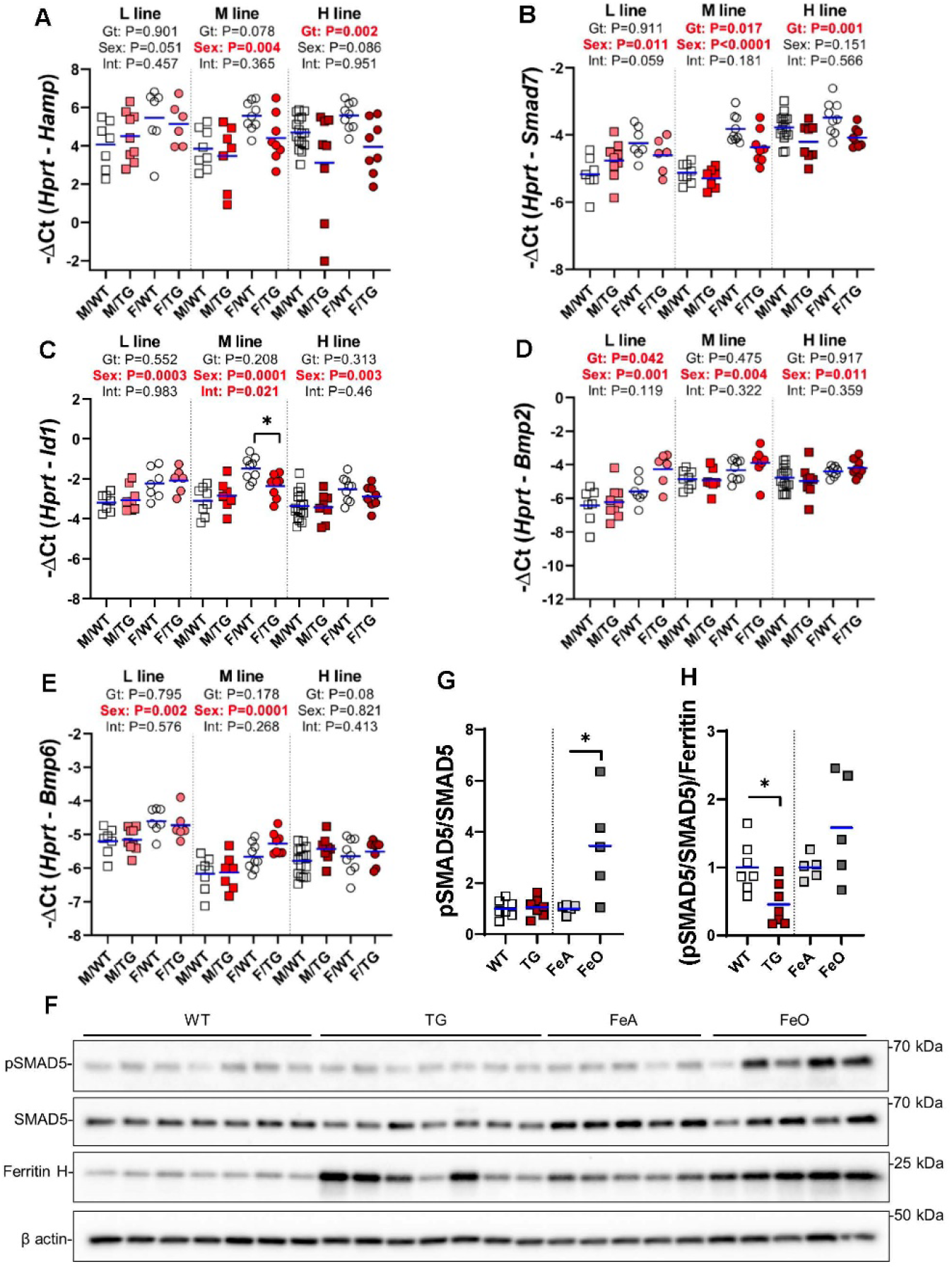
Effect of Erfe overexpression on liver BMP signaling. Relative mRNA expression of **A)** Hamp, **B)** Smad7, **C)** Id1, **D)** Bmp2 and **E)** Bmp6 in the liver at 6 weeks of age in male (M, square) and female (F, circle) wild-type (wt, white symbols) Erfe-overexpressing (TG, colored symbols) mice and wild-type (WT) littermate controls from line-L (white/pink), line-M (white/red), and line-H (white/dark red). Group means are indicated by blue lines and groups within each individual line and age group were compared by two-way ANOVA to determine significant effects of genotype and sex on data variation and to identify interactions between these variables (P<0.05 denoted in bold red). In the event of significant interaction between genotype and sex, individual groups were compared by Šidak’s multiple comparisons test (NS=P>0.05, *=P<0.05, **=P<0.01, ***=P<0.001, ****=P<0.0001). **F)** Western blotting of liver total cell lysates from 6-week-old, male, WT or TG littermates and 16-week-old, male, WT mice fed either an iron-adequate (FeA, light grey symbols) or iron-loaded (FeO, dark grey symbols) diet for pSMAD5, SMAD5, ferritin H, and β actin. Densitometry analysis of **G)** pSMAD5 levels normalized to SMAD5 and **H)** the pSMAD5/SMAD5 ratio normalized to ferritin H levels. Differences in group means between WT and TG mice or FeA and FeO mice, respectfully, were analyzed for statistical significance by Student’s t-test (*=P<0.05).

We further assessed liver SMAD5 phosphorylation in line-H mice and found it similar to that of their WT littermates, but lower compared to WT mice fed iron-rich diet to match their hepatic iron content **(Figure 4F,G, supplemental Figure 3)**. Indeed, when normalized to the expression of liver ferritin, a marker of liver iron stores, the ratio pSMAD5 to total SMAD5 was significantly lower in line-H mice (**Figure 4H)**, suggesting that ERFE blunts BMP signaling caused by iron loading.

In the kidney **(supplemental Figure 4)**, expression of *Hamp* and the sensitive BMP reporter *Id4^26^* were significantly lower in the line-H mice compared to WT, with a similar trend in *Id1*. Expression of *Smad7*, *Bmp2* and *Bmp6* were not different between WT and *Erfe*-overexpressing mice. Compared to the WT littermates, we detected no effect of *Erfe* overexpression on kidney SMAD5 phosphorylation.

We also analyzed the effect of transgenic erythroid *Erfe* in the fetal liver, where erythropoiesis occurs in close proximity to hepatocytes. In e18.5 embryos from all lines, hepatic *Hamp* expression was markedly suppressed in *Erfe*-overexpressing mice compared to WT, in a dose-dependent fashion **(supplemental Figure 5)**. *Id1* expression was lower but only in line-H pups. *Smad7* expression and the erythroid marker *Gypa* were not consistently affected by *Erfe*-overexpression in the fetal liver. Despite the consistent suppression of *Hamp*, SMAD5 phosphorylation in the fetal liver was resistant to the effect of *Erfe*-overexpression. Only line-H male pups displayed a significantly reduced pSMAD5/SMAD5 ratio that was not statistically significant after normalization to ferritin levels **(supplemental Figure 5F-H)**, suggesting that BMP/SMAD signaling in response to iron functions appropriately in the fetal liver despite excess ERFE production.

The high concentration of erythroid *Erfe*-overexpressing cells in the bone marrow raised the possibility that ERFE may locally suppress BMP signaling^7,11^ in that organ. However, we found no suppressive effect of ERFE on the expression of the BMP target genes *Smad7* or *Id1* at 6 weeks of age in any of the lines, and *Bmp2* or *Bmp6* expression in the bone marrow was also not affected **(supplemental Figure 6A-D)**. We did not detect pSMAD5 by western blotting of marrow cells **(supplemental Figure 6E).**

### Erfe overexpression affects growth and organ size

We next surveyed for effects of ERFE on morphogenesis and homeostasis in potential target tissues. Line-H males had consistently reduced body weights at 3 to 16 weeks, compared with WT littermates **(Figure 5A)**. Line-H females also weighed less than WT until 7 weeks but the difference resolved with age. Line-M mice were also lighter than their WT littermates, again with larger differences in males than females **(supplemental Figure 7A)**. Line-L mice of either sex weighed the same as their WT littermates **(supplemental Figure 8A)**.

**Figure 5:**
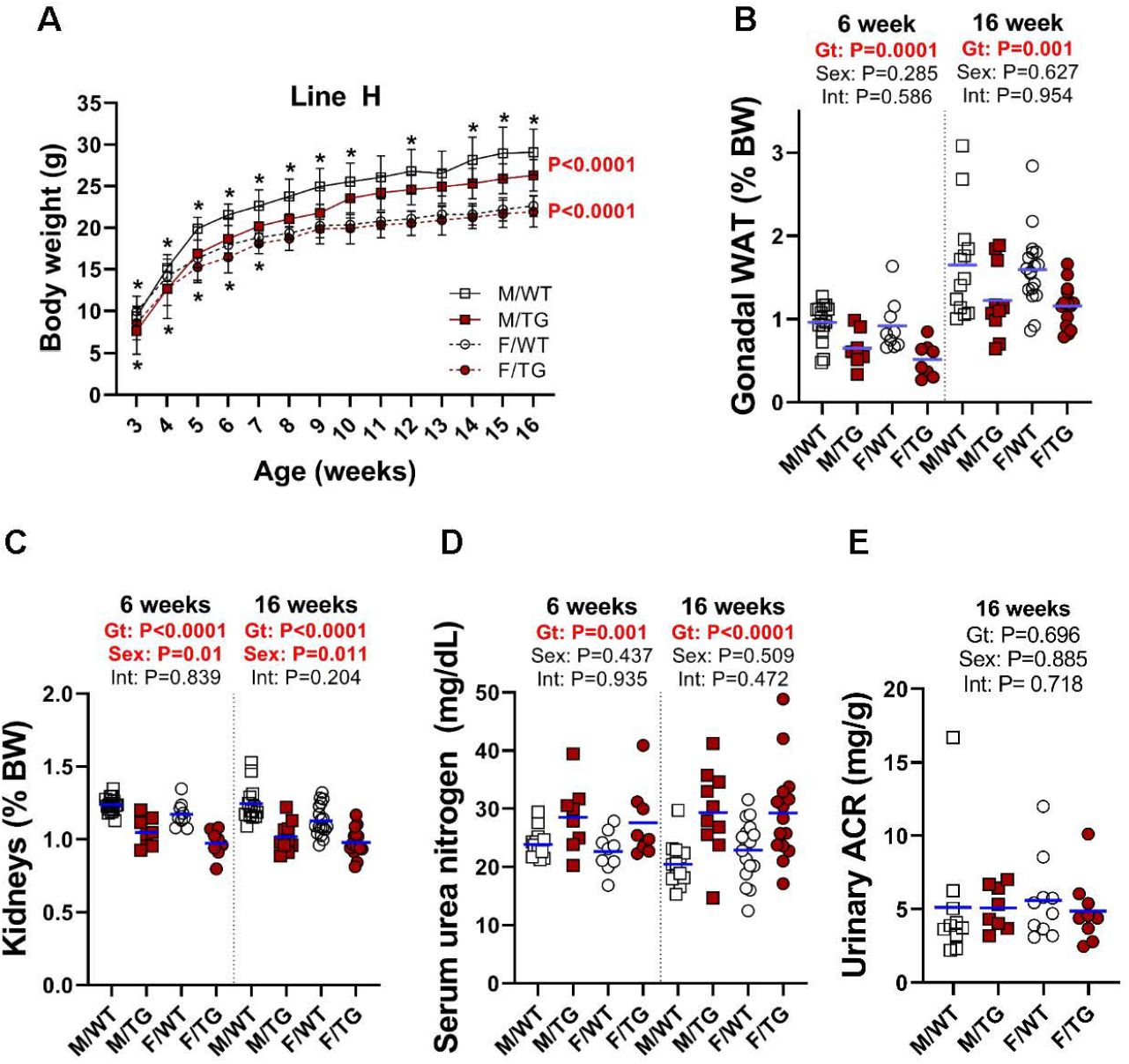
High *Erfe* expression causes reduced body and kidney mass in line-H mice. **A)** Growth curves from ages 3 to 16 weeks of age. Statistical significance of the effect of genotype on data variation between body weights of sex-matched, line-H male (M, square) and female (F, circle) wild-type (WT, white symbols) mice and *Erfe*-overexpressing (TG, dark red symbols) littermates determined by 2-way ANOVA are displayed to the right of their respective curves. Statistically significant differences (P<0.05) between sex-matched WT and TG body weights at individual time points, determined by Student’s *t*-test, are indicated by asterisks (*) above or below curves for males and females, respectively. **B)** Decreased mass of gonadal white adipose tissue (WAT) pads and **C**) kidneys from line-H mice as a percentage of total body weight at 6 and 16 weeks of age. **D**) Serum blood urea nitrogen (BUN) in line-H mice at 6 and 16 weeks of age and **E**) urinary albumin-creatinine ratio (ACR) in line-H mice at 16 weeks of age. Group means are indicated by blue lines. Groups at each age were compared by two-way ANOVA to determine significant effects of genotype and sex on data variation and to identify interactions between these variables (P<0.05 denoted in bold red).

In line-H mice at 6 and 16 weeks, analysis of selected tissues and organs revealed smaller gonadal fat pads and kidneys, even after adjusting for reduced total body weight **(Figure 5B, C)**. Smaller kidney size in line-H mice was associated with reduced kidney function as indicated by elevated serum urea nitrogen levels at 6 and 16 weeks **(Figure 5D)** but we detected no increase in proteinuria in line-H mice compared with WT littermates **(Figure 5E)**. Serum creatinine was also elevated in line-H mice at 16 weeks of age but kidney morphology did not appear altered, and kidney iron status, reflected by renal *Tfrc* expression, did not differ between WT and *Erfe*-overexpressing mice **(supplemental Figure 9)**. Line-H mice at 6 weeks also had significantly reduced inguinal fat pads and increased spleen weights, relative to total body weights, but these differences were not observed at 16 weeks. Brain weights **(supplemental Figure 10)** from transgenic mice comprised a greater relative proportion of total body weight compared with those of WT littermates but this was primarily attributable to the lower body weights of transgenic mice rather than an absolute difference in brain weights.

In line-M mice, kidney mass trended lower compared with WT littermates but this was borderline significant at 6 weeks and significant at 16 weeks for females only. 6-week transgenic females had smaller gonadal and inguinal fat pads. No other differences in individual tissue weights were detected **(supplemental Figure 7B-I)**. No effect of *Erfe* overexpression on tissue or organ weights was detected in line-L mice **(supplemental Figure 8B-E)**.

To analyze the effect of erythroid overexpression of *Erfe* on steady-state bone homeostasis (known to be regulated by BMP signaling^27–29^), we performed bone histomorphometry in female line-H mice at 6 weeks. We detected no significant differences in any measured parameters of bone development, mineralization, or cell populations **(supplemental Table 2)**. Because H-line mice show increased physical activity (see below), we examined mRNA expression of bone-associated markers *Bglap1/2, Sp7 and Runx2* in line-M female mice but detected no significant differences compared with WT (**supplemental Figure 11**).

### Erfe overexpression impairs pup survival and causes developmental and behavioral abnormalities

Unexpectedly, fewer line-H mice survived to weaning than their WT littermates, independently of dam genotype **(Figure 6A)**. To determine if this divergence from expected mendelian inheritance was attributable to loss of pups during embryonic development, we analyzed genotypes from line-H pups between embryonic day 17.5 and 18.5. The proportion of transgenic to WT embryos did not significantly differ from the expected ratio of 50% (**Figure 6B)**. Thus, decreased survival of H-line transgenic mice to weaning is a result of higher perinatal or postnatal mortality. Line-H transgenic mice also occasionally presented with permanent eyelid closure of a single eye (about 5% incidence). Unilateral anophthalmia was seen in developing embryos between days E17.5 and 18.5 in approximately 5% of transgenic mice but not in WT littermates **(Figure 6C)**.

**Figure 6:**
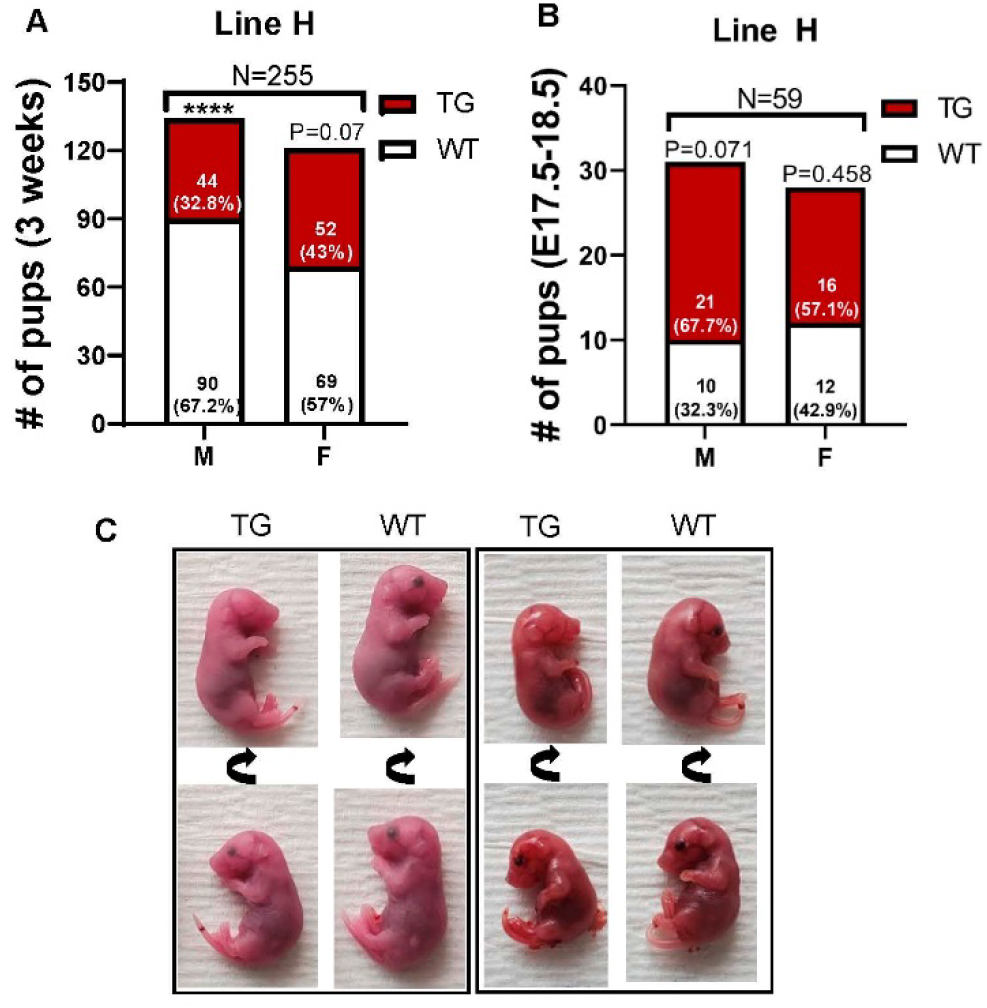
Impaired postnatal survival and unilateral anophthalmia in line-H transgenic mice. Number and within-sex percentage of WT (WT, white) and *Erfe*-overexpressing (TG, dark red) offspring, grouped by offspring sex (male=M, female=F), from line-H breeding surviving at **A)** 3 weeks of age or **B**) in utero at embryonic day e17.5-18.5. Statistically significant differences from expected proportions were assessed by binomial testing (****=P<0.0001). **C**) Examples of unilateral anophthalmia in line-H TG mice at day e17.5-18.5. Top and bottom panels show the same pup rotated to demonstrate, in TG mice, normal (bottom) or impaired (top) eye development compared with WT littermates.

No divergence from expected transgene inheritance ratios was observed in litters from either line-L or line-M breeding (**supplemental Figure 12)** nor any eye abnormalities.

Line-H mice (but not lines-M or -L) displayed compulsive circling behavior detectable as early as 3 weeks and persistent with aging **(Supplemental video file),** affecting 38% of line-H mice **(supplemental Figure 13)**. The H-line mice were more active as indicated by distance run during open field testing. Compared with WT littermates a significantly higher percentage of line-H mice displayed impairment in either righting or negative geotaxis, suggesting that high *Erfe* overexpression is associated with altered vestibular-motor function. *Erfe*-overexpressing and WT mice were equally responsive to auditory stimulation and demonstrated similar physical capacity during a wire maneuver test (per behavioral report).

## Discussion

We present evidence that constant, high-level *Erfe* expression by erythroid cells is sufficient to attenuate hepcidin responsiveness to iron loading and to cause systemic iron overload, manifested by increased plasma iron concentrations and hepatic tissue iron stores. Serum ERFE levels in the 3 lines of transgenic mice ranged from ~0.5 ng/ml, similar to levels we measured contemporaneously in the thalassemia intermedia model Hbb^th3/+^ mice, to ~200 ng/ml. Differences in standards between mouse and human assays and potential differences in the relative potency of mouse and human erythroferrone preclude direct comparison of ERFE concentrations between mouse models and humans. Nevertheless, the broad range of serum ERFE concentrations in our transgenic mice likely encompasses the broad range of ERFE levels reported in humans, including those with hemolytic anemias with ineffective erythropoiesis^12,20^. In our mouse models, the large changes in ERFE protein concentrations that result from much smaller changes in *Erfe* mRNA expression suggest decreasing clearance of plasma ERFE at high concentrations. Comparing liver iron content between the 3 transgenic lines indicates that the effect of erythron-derived ERFE on iron loading is dosedependent but nonlinear.

Liver hepcidin expression is feedback-regulated by liver iron content and serum iron concentrations^11,24^. Any suppression of hepcidin by ERFE will cause increased duodenal iron absorption and hepatic iron accumulation, in turn raising hepcidin concentrations and inhibiting iron absorption, eventually reaching hepcidin concentrations that maintain iron balance. Iron loading in *Erfe*-overexpressing mice changed little between 6 and 16 weeks, indicating that by the age of 6 weeks iron balance was reached, a time course similar to that seen in Hbb^th3/+^mice^30^. The delay in establishment of the iron/hepcidin balance in line-H mice indicates that higher circulating levels of ERFE require greater accumulation of iron to stimulate hepcidin expression sufficiently to overcome the effects of ERFE. In all transgenic lines, hepcidin levels were inadequate when considered in the context of serum and hepatic iron concentrations.

Line-H *Erfe*-overexpressing mice had elevated blood hemoglobin concentrations at 6 and 16 weeks caused by higher erythrocyte hemoglobin content when compared with WT littermates. Line-M mice also had increased hemoglobin concentrations at 16 weeks but not at 6 weeks when rapid growth stresses the iron supply. These effects are consistent with ERFE-dose-dependent hepcidin suppression and consequently enhanced iron availability, as hemoglobin synthesis in mice and humans is modestly enhanced by hepcidin suppression and increased iron availability when erythroid precursors are normal ^31,32^. On the other hand, with pathological erythropoiesis such as in the mouse model of β-thalassemia, hepcidin suppression and increased iron supply <u>worsen</u> ineffective erythropoiesis and anemia, as shown by improved hemoglobin concentrations after interventions that increase hepcidin ^33,34^. A recent report of the detrimental effects of a genetic *ERFE* variant that increased ERFE production in congenital dyserythropoietic anemia type II supports the possibility that chronically elevated ERFE production and the resulting excess of iron may also worsen anemia in human disorders of ineffective erythropoiesis ^35^.

Because the proposed mechanism by which ERFE suppresses hepcidin is BMP sequestration ^7,8,11^, we examined our model for evidence that the effects of ERFE were mediated by suppression of BMP signaling. In addition to *Hamp*, we assessed the expression of the BMP-responsive genes *Id1*^36^ and *Smad7*^37^ in the liver, a known target tissue for ERFE, as well as in the fetal liver and bone marrow, two organs that could be subject to the autocrine or paracrine effects of erythroid-derived ERFE. In the kidney, an organ whose growth was impaired in the highest ERFE model, we also assayed the sensitive BMP signaling marker *Id4^26^*. *Erfe*-overexpressing mice had altered expression of some BMP target genes, but not in all of the organs examined. In the liver, we identified a suppressive effect of moderate or high *Erfe* overexpression on *Hamp* and *Smad7*, but not *Id1* expression. We also found that *Erfe*-overexpressing mice had a blunted hepatic SMAD5 phosphorylation response to iron overload when compared to similarly iron-loaded WT mice. In the fetal liver, we detected suppression of *Hamp* and *Id1*, and in the kidney of *Hamp* and *Id4* but did not detect any corresponding differences in SMAD5 phosphorylation. We did not detect a decrease in the expression of *Id1* or *Smad7* in the bone marrow of *Erfe*-overexpressing mice compared with WT littermates, nor any compensatory increase in *Bmp2* and *Bmp6* expression by cells in the marrow. Differential sensitivity of tissues to the effects of ERFE suggests that the mechanism of ERFE action may depend on differences in local BMP-signaling mechanisms, perhaps mediated by tissue-specific coreceptors^38,39^. Our findings also leave room for the possibility that the current model of the mechanism of ERFE effect on hepcidin expression is incomplete.

While the abnormalities in transgenic mice with low circulating levels of ERFE were entirely limited to iron homeostasis, line-H mice with high ERFE levels displayed increased postnatal mortality, reduced body weight and gonadal adipose tissue weight, reduced kidney size and function, occasional unilateral anophthalmia, and compulsive circling behavior. In a weaker echo of these abnormalities, the moderately *Erfe*-overexpressing line-M mice also showed lower body mass and smaller kidneys relative to body mass than their WT littermates. We considered the possibility that these abnormalities were caused by disruption of a critical gene at the site of transgene insertion or interference with topologically-associating domains. We found no evidence for this possibility by genomic sequencing, which confirmed that line-H is derived from the same founder as line-M and shares the same insertion site on Chr. 4, overlapping a gene for a noncoding RNA of unknown function. The lines H and M only differ in transgene copy number, with higher copies resulting in increased phenotypic severity. Furthermore, the corresponding genomic sequence on the other copy of Chr. 4 is WT. Rather, the spectrum of abnormalities is consistent with disruption of signaling by select BMPs during embryonic development or early growth. BMP signaling plays a key role in a wide range of physiological processes involved in both the development and homeostatic maintenance of various organs ^29,40–42^. ERFE was also identified as a vascular morphogen in Xenopus embryos where it acts by antagonizing BMP4 ^43^. Although direct application of the mouse or Xenopus models to human disease is too simplistic, our observations raise the possibility that some of the nonhematologic complications experienced by patients with ineffective erythropoiesis ^14–16^ could either be caused or exacerbated by disrupted BMP signaling.

Specifically, deletion of *Bmp7* in mice leads to impaired embryonic kidney development ^44,45^. Eye development is also regulated, in part, by BMP signaling ^40,44^ and disruption of BMP signaling by conditional deletion of BMP receptors in the eye during embryonic development leads to anophthalmia ^46^. BMP4 is required for normal vestibular function and BMP4^+/-^ mice demonstrate circling behavior ^42^, similar to line-H mice in the present study. While mouse ERFE is reported to not inhibit BMP4 ^43^, ERFE may interact with and inhibit BMP4 heterodimers formed with other BMPs, e.g. BMP7 ^47^. Adipogenesis is regulated by nuanced interplay between BMP signaling and antagonism ^48,49^. A preferential effect of ERFE on visceral fat compared with subcutaneous fat, as the present study suggests, may be due to differences in the expression of, or receptivity to, specific BMPs by adipocytes or adipose progenitor cells in these compartments ^50 51^. ERFE is reported to alter BMP2 signaling in a “depot-specific” manner *in vitro*, altering BMP signaling to a greater degree in adipocytes derived from some adipose compartments compared with others ^52^. Reduced adiposity in response to *Erfe* overexpression is consistent with previous findings of increased adipose deposit size in *Erfe*-deficient mice ^53^.

Altered postnatal survival and decreased body weights were more prominent in male line-H mice compared with females, despite similar circulating ERFE levels between sexes. We also noted a strong pattern of increased hepatic and renal BMP2 and BMP6 expression in female compared with male mice of all genotypes (Figures 4DE, supplemental figure 4EF). Such effects of sex on BMP2 and 6 expression were reported to be driven by estrogen signaling ^54,55^. Increased production of BMP2 and BMP6 in females could compensate for BMP inhibition by ERFE.

A well-documented complication of β-thalassemia is a loss of bone mineral density associated with marrow expansion driven by chronic erythropoietic stimulation ^56^. We observed no clear difference in steady-state bone histomorphometry between line-H mice and WT controls or in expression of bone-associated markers in line-M compared to WT controls. The lack of effect of ERFE on bone development and mineralization was unexpected given the substantial effects, including spontaneous fractures, observed in bones of mice overexpressing other BMP antagonists ^57,58^. However, in these studies BMP antagonists were overexpressed selectively in cells of the osteoblast lineage and likely exhibited autocrine inhibition of BMP signaling in osteoblasts, unlike in the present study where endocrine or paracrine BMP inhibition by ERFE predominated. In β-thalassemia mice, ERFE was recently reported to have an osteoprotective effect, but this phenotype may also be dependent on osteoblast ERFE expression^59^ not examined in our model.

Kidney dysfunction in the context of β-thalassemia is incompletely understood but thought to result from a combination of iron-mediated damage, chronic anemia with renal hypoxia, and treatment with iron chelators that may damage the kidney ^60^. Signaling by BMP7 is thought to protect against renal fibrosis ^61^ and increased expression of BMP antagonists may predispose the kidney to tubular injury ^62^, exacerbating damage from excess iron or chelation therapy. Reducing the amount or activity of ERFE in patients with β-thalassemia may represent a potential therapeutic option for preventing kidney damage. It remains to be determined which aspects of the phenotype observed in mice with high circulating ERFE levels in this study are the result of altered BMP signaling in utero versus during postnatal life. Relative to human β-thalassemia, transgenic mice in this study may be more susceptible to abnormal organ development because of differences in the timing of increased *Erfe* expression during embryonic and postnatal development. In humans with β-thalassemia, anemia and ineffective erythropoiesis does not develop until after birth ^63^ because of the protective effect of fetal hemoglobin, whereas transgene expression in our mouse model begins during embryonic development^22^. Therefore, increased *Erfe* expression, resulting from anemia in patients with β-thalassemia, is not expected until the postnatal period, diminishing the potential effects on organ development. However, elevated ERFE levels in utero would be expected in patients with fetal anemias such as α-thalassemia ^64^, pyruvate kinase deficiency ^65^, congenital dyserythropoietic anemias ^66^ and other congenital anemias with ineffective erythropoiesis, and could contribute to developmental abnormalities seen in their severe forms ^67^.

In conclusion, our mouse models demonstrate that chronically increased blood ERFE levels contribute to iron loading and, at higher levels, alter the development or homeostasis of multiple organ systems. ERFE may be a suitable therapeutic target in anemias from ineffective erythropoiesis to reduce iron loading and possible organ dysfunction from ERFE-mediated inhibition of BMP signaling.

## Acknowledgments

Support for this work was provided by NIH R01DK126680 (TG) and Cooley’s Anemia Foundation (RC). The authors thank the UCLA Translational Pathology Core Laboratory for histology processing, the UCLA Technology Center for Genomics & Bioinformatics (TCGB) for performing genomic sequencing, and the UCLA Behavioral Testing Core for performing mouse behavioral analysis.

## Author Contributions

RC designed and performed experiments, analyzed data, and wrote the manuscript. GJ, JDO and GK performed experiments and assisted with data interpretation. RCP performed bone analyses and assisted with their interpretation. EN and TG conceived the project, designed experiments, analyzed data, and wrote the manuscript.

