## Supplemental Materials, Tables and Figures for "Erythroid overproduction of erythroferrone causes iron overload and developmental abnormalities in mice"

**Materials and Methods**

*Animal husbandry and diet*

Unless otherwise specified, animals were fed a natural ingredient diet (Labdiet 5053) containing ~220 ppm iron. Animal body weights were recorded weekly beginning at 3 weeks and continuing until euthanasia, either at 6 or 16 weeks of age. In experiments aimed at altering systemic iron levels, C57BL/6J mice were purchased from The Jackson Laboratory and were fed either an iron-adequate or iron-loaded purified-ingredient diet (Envigo) based on the AIN-76A formulation containing either 48 ppm ferric citrate or 5,000 ppm carbonyl iron, respectively, between the ages of 13 and 16 weeks. *Hbb^th3/+^* mice used for the comparison of serum ERFE values with *Erfe*-overexpressing mice were generously contributed by Dr. Stefano Rivella. All animals were housed in a specific-pathogen-free barrier facility. Experimental animals were euthanized by isoflurane inhalation. All experiments involving mice were conducted with the approval of the University of California, Los Angeles Animal Research Committee.

*Genomic characterization*

Genomic DNA was prepared from the livers of the H-line transgenic founder, H-line F1 progeny, M-line founder, and one M-line F1 progeny using the Gentra Purgene Mouse Tail Kit (Qiagen) per manufacturer’s instructions. Genomic sequencing on paired-end libraries was performed at UCLA Technology Center for Genomics & Bioinformatics using a Novaseq S4 DNA sequencer (Illumina). The reads were aligned to the reference genomic sequence Mus musculus strain C57BL/6J chromosome 4, GRCm38.p6 C57BL/6J and to the transgene sequence, and overlap sequences between adjacent transgene copies and between transgene and chromosomal sequence identified and used to ascertain the chromosomal insertion site, and transgene copy number.

*Measurement of iron-related and hematological parameters*

Serum iron and tissue non-heme iron were measured by using a colorimetric assay according to the manufacturer’s protocol (Sekisui Diagnostics). Prior to sampling, tissues were homogenized to reduce variation resulting from regional differences in iron deposition. Hemoglobin levels, red-blood-cell counts, and mean corpuscular hemoglobin levels were determined by using a HemaVet blood analyzed (Drew Scientific). Serum ferritin concentrations were measured by the Mouse Ferritin ELISA Kit (Immunology Consultants Laboratory, Inc.) and serum transferrin concentrations were measured by using the Mouse Transferrin ELISA Kit (Alpha Diagnostic International), following the manufacturer’s protocols. Transferrin saturation was calculated by measuring serum iron and serum transferrin in each sample and determining the ratio of serum iron to total transferrin iron-binding sites on a molar basis.

*Quantification of serum and urinary proteins and metabolites*

Serum ERFE concentrations were determined as previously described ^1^ with the substitution of DELFIA europium-conjugated streptavidin for horseradish peroxidase (HRP) conjugated streptavidin or with a commercially available kit HRP-based, anti-ERFE ELISA kit sharing the same ERFE standard (Intrinsic LifeSciences). Fluorescence was measured by CLARIOstar Plus microplate reader (BMG Labtech). Serum hepcidin concentrations were determined by ELISA as detailed previously ^2^. Serum urea nitrogen concentrations were measured by using the QuantiChrom Urea Assay Kit (BioAssay Systems), urinary albumin using the Mouse Albumin ELISA Kit (Abcam, ab108792), urinary creatinine using the Creatinine (urinary) Colorimetric Assay Kit (Cayman Chemical), and serum creatinine using the Creatinine (serum) Colorimetric Assay Kit (Cayman Chemical), all in accordance with the manufacturers’ protocols.

*RNA isolation and measurement of gene expression*

Total RNA isolation was performed by using TRIzol (ThermoFisher Scientfic). cDNA was synthesized by using the iScript cDNA Sythesis Kit (Bio-Rad) following the manufacturer’s protocol. Relative mRNA expression for genes of interest were determined by quantitative RT-PCR using SsoAdvanced Universal SYBR Green Supermix (Bio-Rad) and measured on a CFX-96 RT-PCR Detection System (Bio-Rad). Primer sequences used in this study are listed in Supplemental Table 1.

*Behavioral testing*

All behavioral testing was performed at the UCLA behavioral testing core facility by personnel blinded to the mouse genotype. Mice were transferred to the testing facility and allowed 1 week to acclimate prior to testing between 11 and 13 weeks of age. Open-field testing was performed for a duration of 30 minutes in an empty and dimly lit, light intensity less than 20 lux, square area measuring 30 cm per side. Mouse movement was recorded automatically and analyzed by using ANY-maze software (Stoelting). Righting reflex of experimental animals was assessed by flipping animals held by the tail, so that the animal passed through an inverted position, to an approximate height of 25 cm. Landing position of the mouse was recorded and a score of 0, 1, or 2 given to mice that landed on their feet (0), side (1), or back (2), respectively. Righting reflex testing was performed three times per mouse and the highest value recorded. Negative geotaxis testing was assessed by placing mice facing downwards on a declined surface. Mouse movement was recorded for a duration of 30 s and a score of 0, 1, 2, or 3 was given to mice that turned to face upslope and climbed (0), turned without climbing (1), moved without turning (2), or did not move (3). Reaction to auditory stimulation was measured by the degree of visible response to a 20 KHz sound with a volume of 90 dB at mouse location. A score of 0, 1, 2, or 3 was given to mice that did not respond (0), exhibited a pinna reflex (1), jumped < 1 cm (2), or jumped > 1 cm (3). Wire maneuvering capacity was assessed by the ability of mice to hang onto a wire and navigate to an attachment post when allowed to initially establish a foreleg grip. A score of 0, 1, 2, 3, or 4 was assigned to mice that demonstrated active hindleg grip (0), impaired hindleg grip (1), inability to grip with hindlegs (2), inability to lift legs followed by falling (3), or immediately falling (4).

*Bone histomorphometry analysis*

Mouse femurs were dehydrated and fixed in 70% ethanol prior to embedding in methyl methacrylate. Sections cut longitudinally at a thickness of 5 µm were stained with toluidine blue (pH 6.4) and used for static bone parameter analysis on a total of 20 fields per sample by using a OsteoMeasure morphometry system (Osteometics) at a distance of 200 µm from the growth plate. Results for individual parameters are listed using accepted units and terminology recommended by the Histomorphometry Nomenclature Committee of the American Society for Bone and Mineral Research ^3^.

*Tissue histology analysis*

Tissue samples were fixed in 10% neutral buffered formalin for 24 h followed by 70% ethanol storage prior to paraffin processing and embedding. Sections were cut at a thickness of 4 µm and stained on an automated device with hematoxylin and eosin for morphological analysis. Staining for tissue iron deposition was performed using enhanced Perls’ Prussian blue staining. In brief, tissue sections were deparaffinized and rehydrated followed by incubating with a mixture of 2% potassium ferrocyanide and 4% hydrochloric acid for 30 minutes. After washing with tap water, samples were incubated for 20 minutes with 0.3% hydrogen peroxide, to quench endogenous peroxidase activity, washed again in tap water and incubated with ImmPACT® SG enhancement reagents (Vector Laboratories, SK-4705) following the manufacturer’s protocol. Sections were briefly counterstained with hematoxylin solution, Gill #1 (Sigma-Aldrich) prior to dehydration and mounting. Images were captured using an Eclipse E600 microscope (Nikon) with SPOT Basic^TM^ (SPOT Imaging) image capture software.

*Immunoblotting*

Tissues were homogenized in RIPA lysis buffer (Santa Cruz) including 1x Halt^TM^ Protease and Phosphatase Inhibitor Cocktail (ThermoFisher Scientific). Lysate protein concentrations were determined by using the Pierce^TM^ BCA Assay Kit (ThermoFisher Scientific). Samples were mixed with a reducing sodium dodecyl sulfate (SDS) sample loading buffer, containing a final concentration of 0.6 M dithiothreitol, and were heated at 95 degrees C for 5 minutes prior to electrophoretic separation on precast 4-20% gradient SDS-polyacrylamide gels (Bio-Rad). Protein was transferred to nitrocellulose membranes and incubated for 1 h in a blocking buffer containing 5% non-fat dry milk dissolved in Tris-buffered saline containing 0.1% Tween 20 (TBS-T). Primary antibodies generated against pSMAD5, 1:2,000 v/v (Abcam, ab92698), SMAD5, 1:2,500 v/v (Abcam, ab40771), ferritin-H, 1:10,000 v/v (Cell Signaling Technology, #4393), β-actin 1:100,000 v/v (Sigma-Aldrich, A3854) were diluted in blocking buffer and incubated with membranes overnight at 4 degrees C. Blots were then washed with TBS-T followed by incubation with an HRP-conjugated donkey anti-rabbit secondary antibody, 1:5,000 v/v (ThermoFisher Scientific, #31462) for 1.5 h at room temperature. Following washing, blots were incubated with either SuperSignal^TM^ West Pico PLUS or West Femto Maximum Sensitivity Substrate (ThermoFisher Scientific) and visualized with a ChemiDoc XRS+ System (Bio-Rad). Densitometric analysis was performed using Image Lab Software (Bio-Rad).

*Statistical analysis*

Statistical analysis was performed by using the statistical package included in Prism version 8 (GraphPad). P<0.05 was considered statistically significant. For each individual transgenic mouse line and its WT littermates, means of experimental groups, defined by genotype and sex, were compared with others by using either student’s *t*-test, Welch’s ANOVA, or Two-way ANOVA where indicated. In the event of significant differences between group means after one-way ANOVA analysis, individual group means were compared by using an unpaired-*t*-test with Welch’s correction. Significant interaction between variables during two-way ANOVA testing was followed by Šidak’s multiple comparisons test to determine significant differences between WT and transgenic mice within each sex. Analysis of observed vs expected ratios when analyzing transgene inheritance was performed by using binomial testing and an expected transgene incidence of 50%. Fisher’s exact test was used to analyze differences in phenotypic incidence between groups.

**Supplemental Tables**

|  | **F:5'---3'** | **R:5'---3'** |
| --- | --- | --- |
| ***mHprt*** | **CTGGTTAAGCAGTACAGCCCCAA** | **CAGGAGGTCCTTTTCACCAGC** |
| ***mErfe*** | **ATGGGGCTGGAGAACAGC** | **TGGCATTGTCCAAGAAGACA** |
| ***mHamp*** | **TTGCGATACCAATGCAGAAGA** | **GATGTGGCTCTAGGCTATGTT** |
| ***mId1*** | **ACCCTGAACGGCGAGATCA** | **TCGTCGGCTGGAACACATG** |
| ***mId4*** | **CTGGAGACTCACCCTGCTTT** | **CTGTCACCCTGCTTGTTCAC** |
| ***mSmad7*** | **TTGCCTCGGACAGCTCAATTC** | **CGCACTTTCTGTACCAGCTGA** |
| ***mBmp2*** | **GATCTGTACCGCAGGCACTC** | **CCGTTTTCCCACTCATCTCT** |
| ***mBmp6*** | **ATGGCAGGACTGGATCATTGC** | **CCATCACAGTAGTTGGCAGCG** |
| ***mGypa*** | **ATGGCAGGGATTATCGGAAC** | **CACCCTCAGGAGATTGGATG** |
| ***mTfrc*** | **TCATGAGGGAAATCAATGATC** | **GCCCCAGAAGATATGTCGGAA** |
| ***mTfr2*** | **CCTACTGCCGCTAGACTTCG** | **CAGAGTACACCCACTGCAGG** |
| ***mTmprss6*** | **GTGGTGTACTCGGCCACTGT** | **GTCCTGAGGGACCAGCTGTA** |
| ***mBglap1/2*** | **AGACAAGTCCCACACAGCAG** | **CTGGGCTTGGCATCTGTGA** |
| ***mSp7*** | **TCTCTCCTGCAGGCAGTCC** | **GGCCCAGGAAATGAGTGAGG** |
| ***mRunx2*** | **CTGAGCCAGATGACATCCCC** | **GTCATCATCTGAAATACGCCTGG** |

**Table S1. Primer sequences used for qRT-PCR**

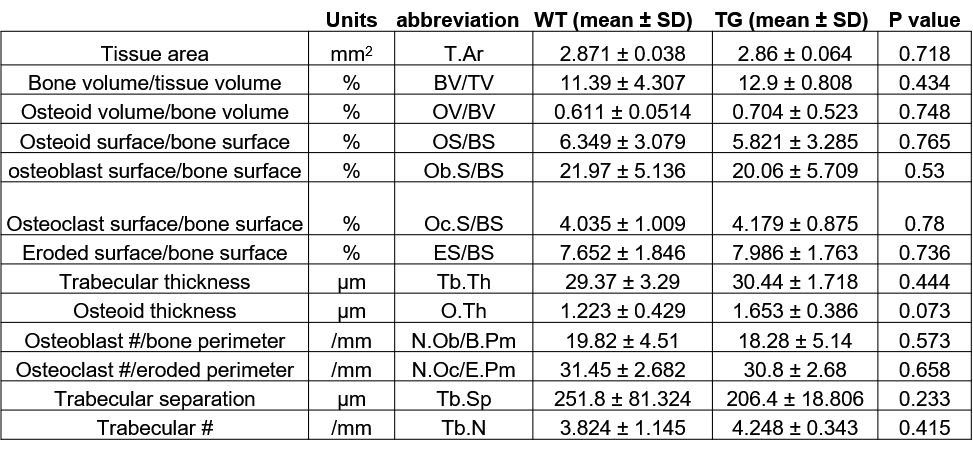

**Table S2.** **Bone histomorphometry analysis.** Parameters obtained from femurs of 6-week-old, female, line-H, wild-type (WT, n=6) and transgenic (TG, n=8) littermate mice were compared using Student’s *t*-test.

**Supplemental Figures**

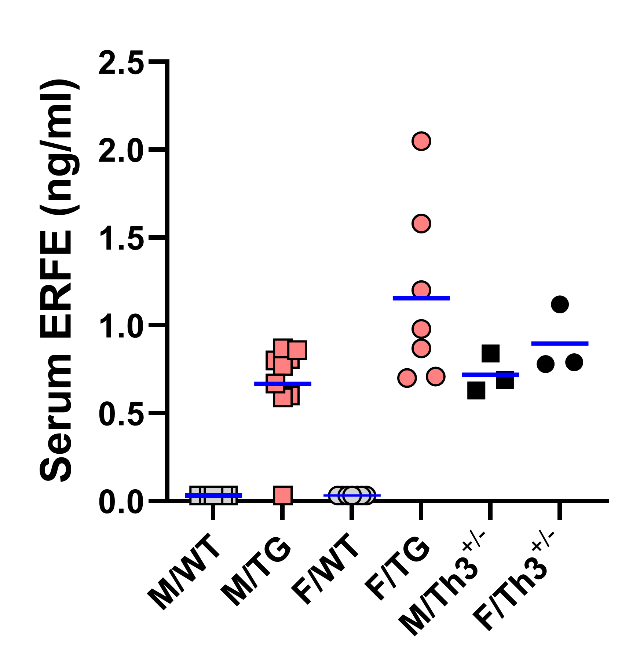
**Supplemental Figure 1**

**Supplemental figure 1. Comparison of serum ERFE levels between Line-L and Hbb^Th3/+^ mice.** Serum ERFE levels in male (M, square) or female (F, circle) Erfe-overexpressing (TG, pink symbols) or wild-type (WT, clear symbols) littermates from line-L or mice heterozygous for the Hbb^Th3^ allele (Th3^+/-^, black symbols) at 16 weeks of age. The mean for each group is indicated by blue line. Within each sex, groups were compared by one-way ANOVA with Tukey’s multiple comparisons testing and groups not sharing a common alphabetical superscript differ significantly (*P*<0.05).

**Supplemental Figure 2**

**A**

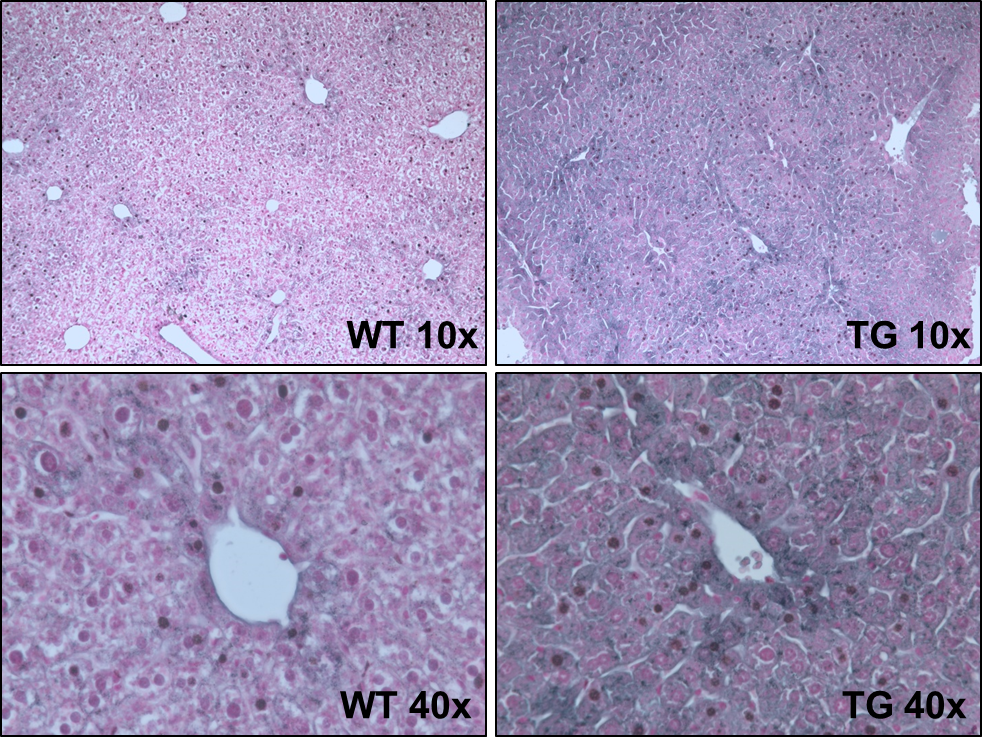

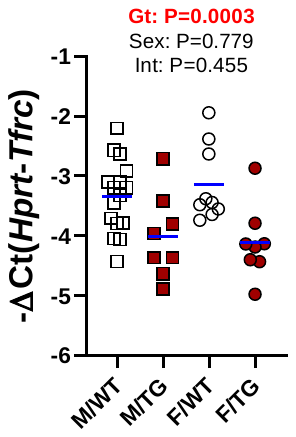

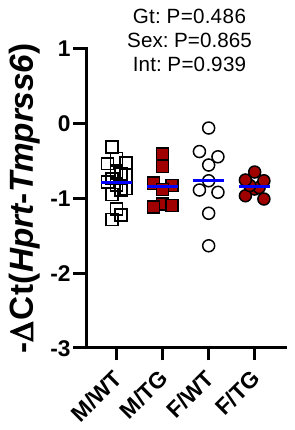

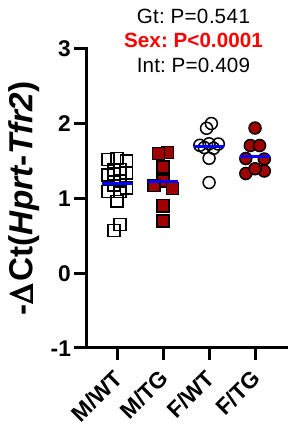

**B**

**C**

**D**

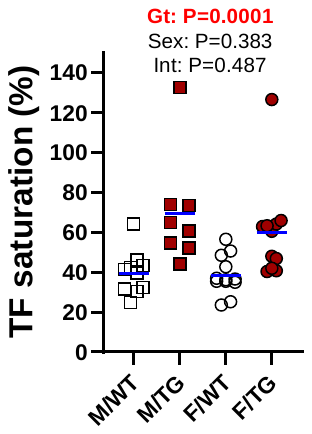

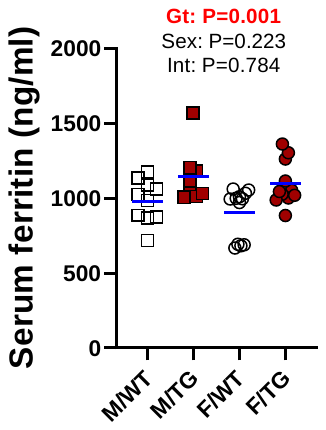

**E**

**F**

**G**

**H**

**I**

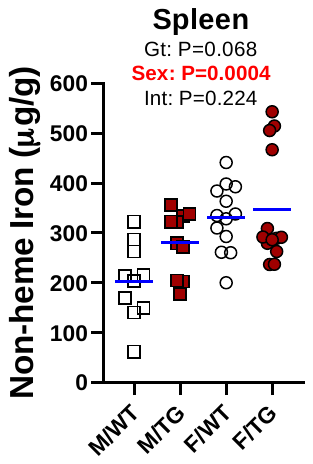

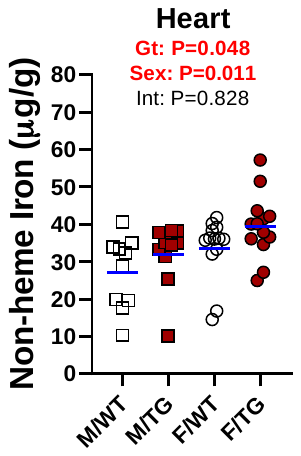

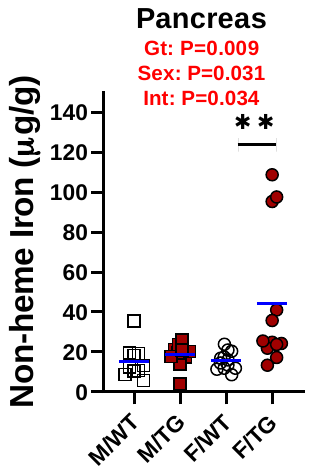

**Supplemental figure 2. Further characterization of iron status in line H mice. A)** Representative DAB-enhanced Perls’ Prussian blue staining of liver sections from wild-type (WT) or *Erfe*-overexpressing (TG) female mice at 16 weeks of age. Iron deposition is indicated by grey/blue staining. Relative liver mRNA expression of **B)** *Tfrc*, **C)** *Tmprss6*, and **D)** *Tfr2* in 6-week-old male (M, square) and female (F, circle) WT (white symbols) and TG (dark red symbols) mice. **E)** Serum ferritin and **F)** transferrin saturation levels in 16-week-old mice. Wet tissue non-heme iron levels in the **G)** spleen, **H)** pancreas, and **I)** heart of line H mice at 16 weeks of age. For panels **B**-**I** group means are indicated by blue lines. Groups were compared by two-way ANOVA to determine effects of genotype and sex on data variation (significant differences denoted in bold red) and to identify interactions between these variables. In the event of significant interaction between genotype and sex, individual groups were compared by Šidak’s multiple comparisons test (ns=P≥0.05, *=P<0.05, **=P<0.01, ***=P<0.001, ****=P<0.0001).

**Supplemental Figure 3**

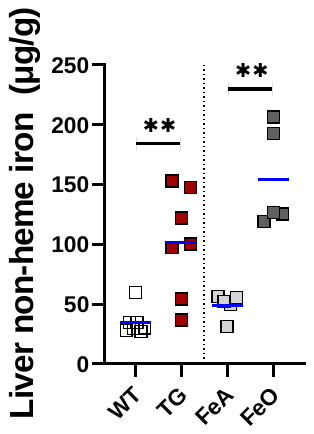

**Supplemental Figure 3. Comparison of liver iron accumulation between genotypes and dietary conditions.** Liver non-heme iron content in male (square) 6-week-old wild-type (WT, white symbols) and *Erfe*-overexpressing (TG, dark-red symbols) or 16-week-old WT mice fed either an iron adequate (FeA, light-grey symbols) or iron-loaded (FeO, dark-grey symbols) diet. Differences in group means between WT versus TG mice and FeA versus FeO mice, respectively, were analyzed for statistical significance by Student’s *t*-test (**=P<0.01).

**Supplemental Figure 4**

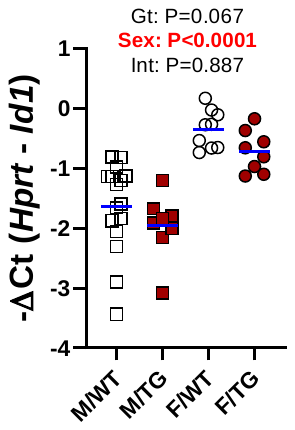

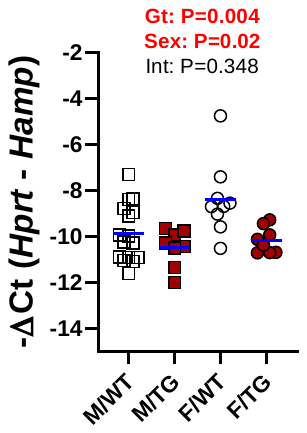

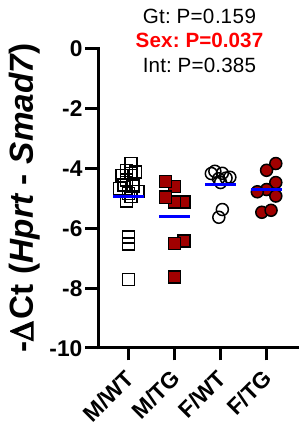

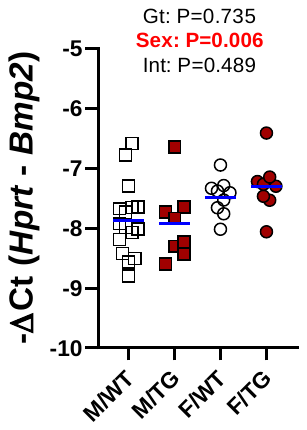

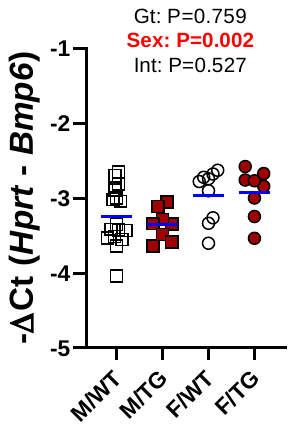

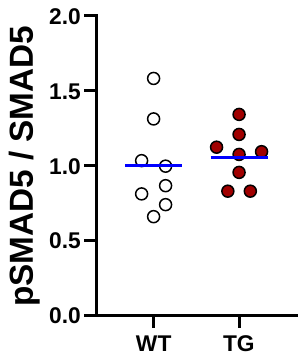

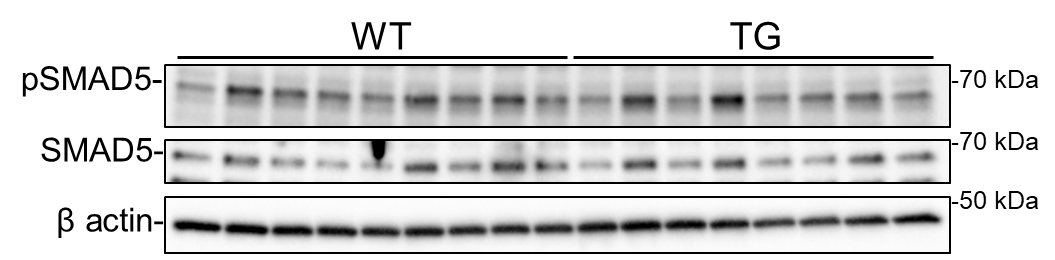

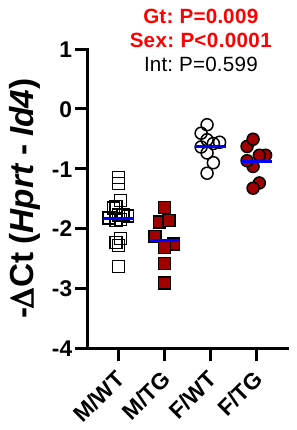

**A**

**B**

**C**

**D**

**E**

**F**

**G**

**H**

**Supplemental figure 4. Effect of *Erfe* overexpression on kidney BMP signaling.** Relative mRNA expression of **A)** *Id1*, **B)** *Id4*, **C)** *Hamp*, **D)** *Smad7* **E)** *Bmp2*, and **F)** *Bmp6* in the kidney at 6 weeks of age in male (M, square) and female (F, circle) wild-type (WT, white symbols) or *Erfe*-overexpressing (TG, dark-red symbols) mice from line-H (white/dark red). Group means are indicated by blue lines and groups within each individual line and age group were compared by two-way ANOVA to determine significant effects of genotype and sex on data variation and to identify interactions between these variables (P<0.05 denoted in bold red). **G)** Western blotting of kidney total cell lysates from 6-week-old, female, WT or TG mice for pSMAD5, SMAD5, and β actin. **H)** Densitometry analysis of pSMAD5 levels normalized to SMAD5 levels. Differences in group means between WT and TG mice were analyzed for statistical significance by Student’s *t*-test (*=P<0.05).

**Supplemental Figure 5**

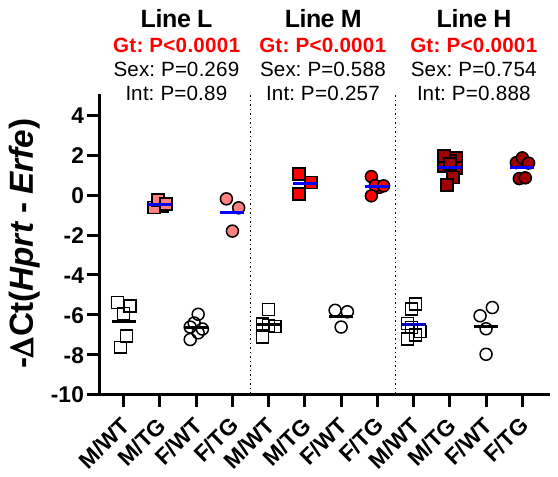

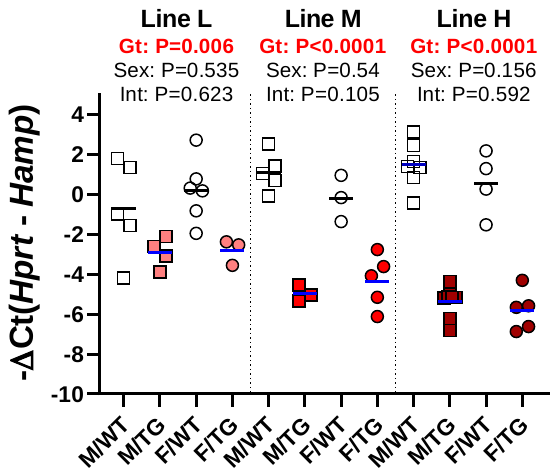

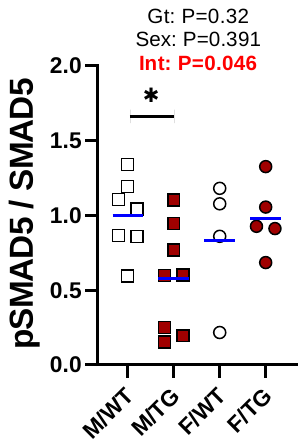

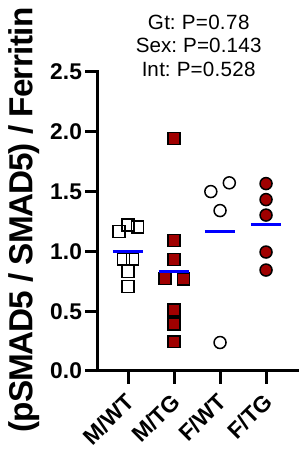

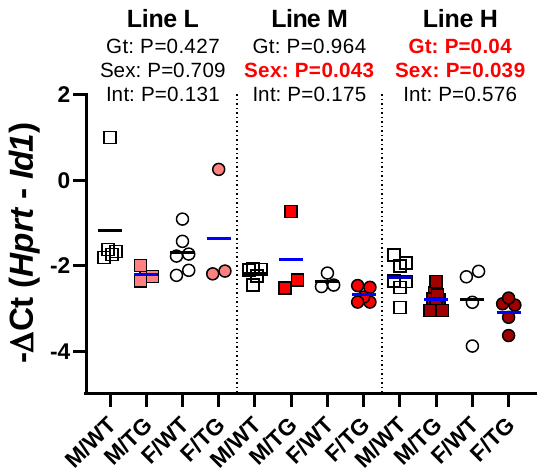

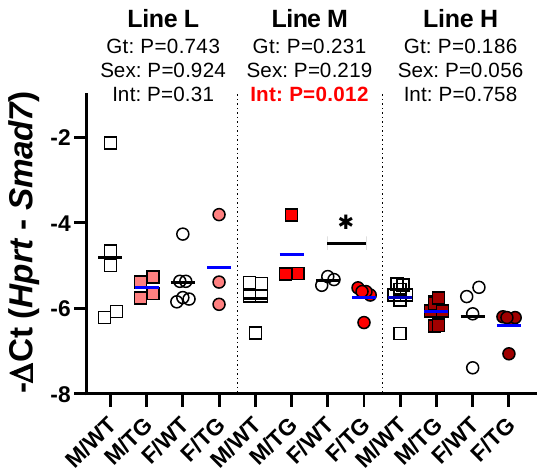

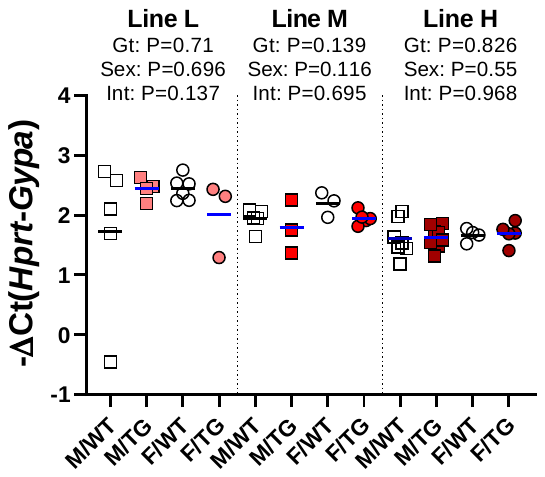

**B**

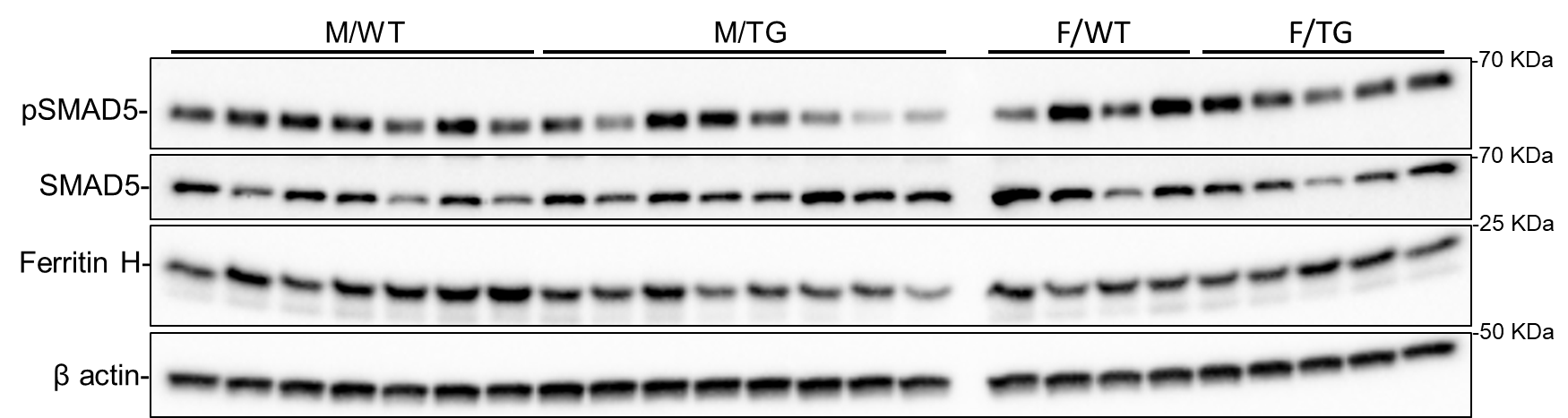

**F**

**H**

**G**

**A**

**D**

**E**

**C**

**Supplemental figure 5. Analysis of fetal liver *Erfe* expression and BMP signaling in *Erfe*-overexpressing mice.** Relative fetal liver mRNA expression of **A)** *Erfe*, **B)** *Hamp*, **C)** *Id1*, **D)** *Smad7*, and **E)** *Gypa* in male (M, square) and female (F, circle) wild-type (WT) mice and *Erfe*-overexpressing (TG) littermates from line-L (white/pink symbols), line-M white/red symbols), and line-H (white/dark red symbols). **F)** Western blotting of fetal liver total cell lysates from line-H mice for pSMAD5, SMAD5, ferritin H, and β actin. Densitometry analysis of **G)** pSMAD5 levels normalized to SMAD5 and **H)** the pSMAD5/SMAD5 ratio normalized to ferritin H levels. Group means are indicated by blue lines and groups within each individual line and age group were compared by two-way ANOVA to determine significant effects of genotype and sex on data variation and to identify interactions between these variables (P<0.05 denoted in bold red) and to identify interactions between these variables. In the event of significant interaction between genotype and sex, individual groups were compared by Šidak’s multiple comparisons test (ns=P≥0.05, *=P<0.05, **=P<0.01, ***=P<0.001, ****=P<0.0001).

**Supplemental Figure 6**

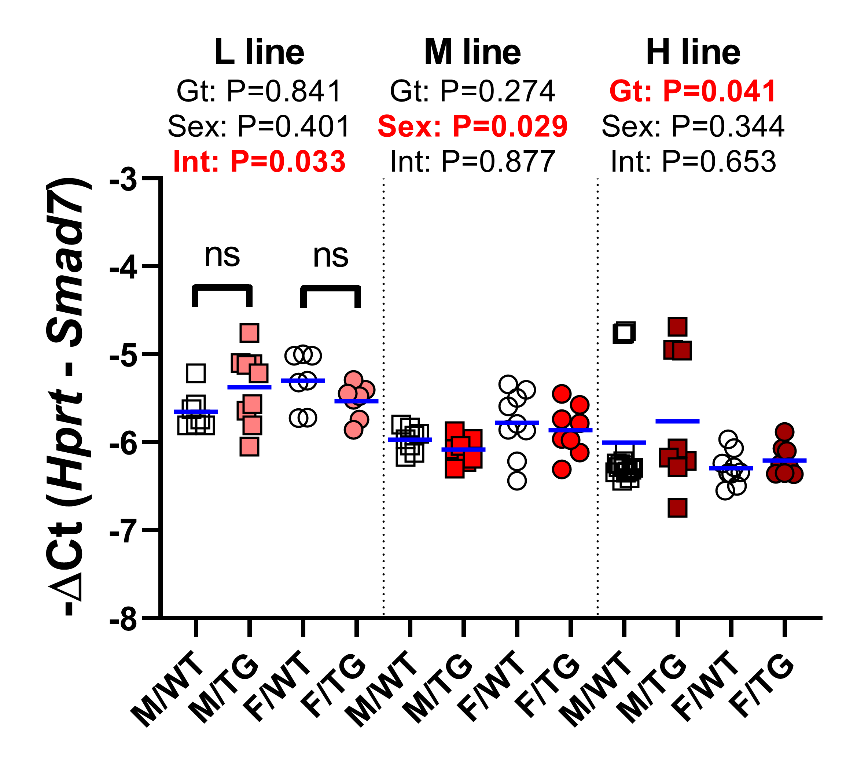

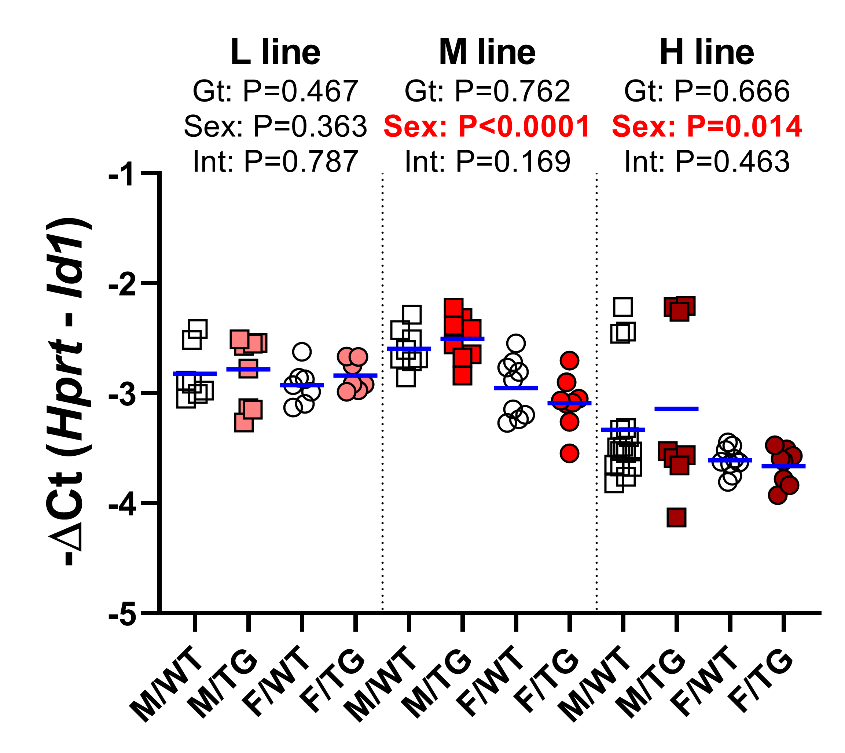

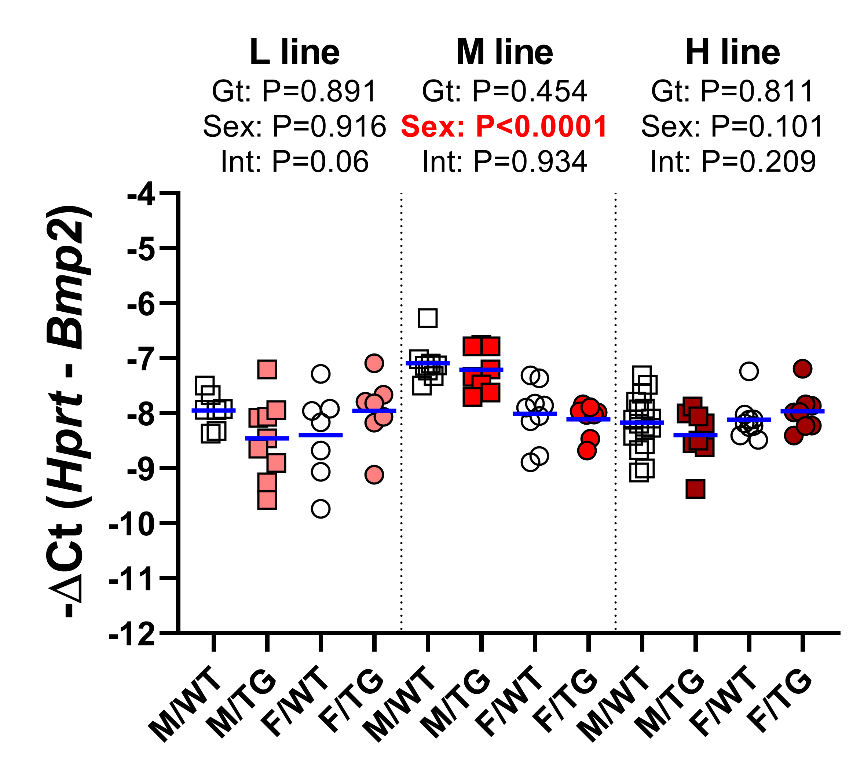

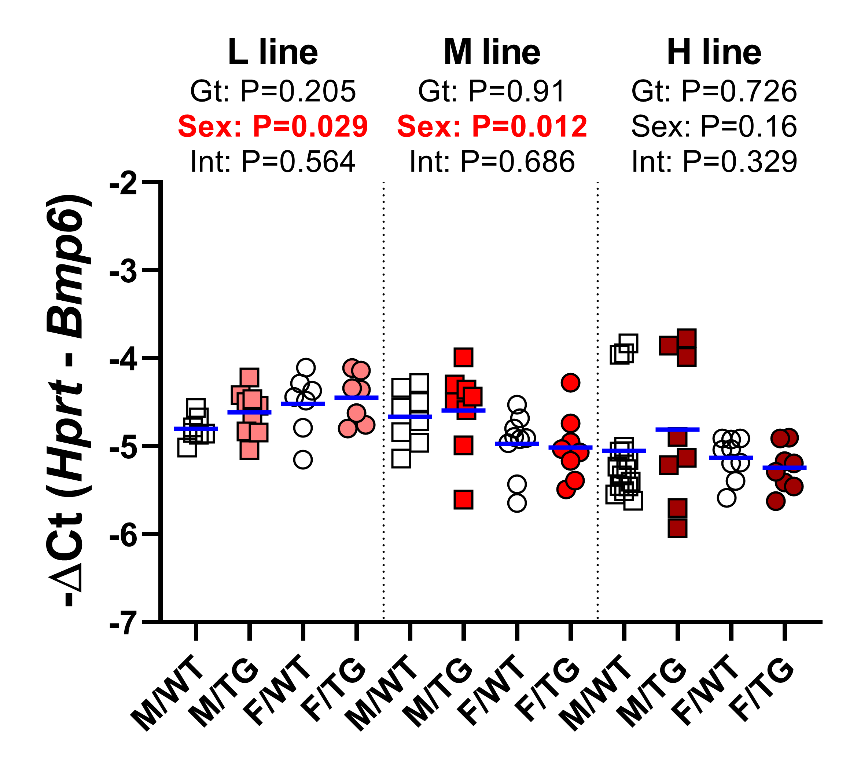

**A**

**B**

**C**

**D**

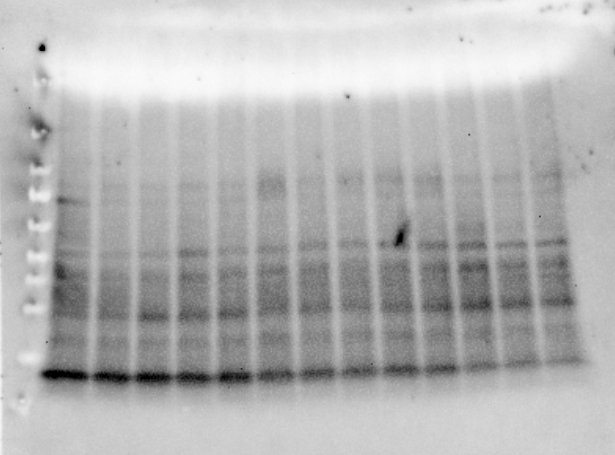

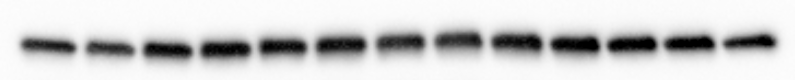

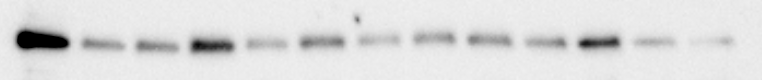

SMAD5-

β actin-

pSMAD5-

-70 kDa

-70 kDa

-50 kDa

WT

TG-M

TG-H

**E**

**Supplemental figure 6. Effect of *Erfe* overexpression on bone marrow BMP signaling.** Relative mRNA expression of **A)** *Smad7*, **B)** *Id1*, **C)** *Bmp2*, **D)** *Bmp6* in the bone marrow at 6 weeks of age in male (M,square) and female (F,circle) wild-type (WT) mice and *Erfe*-overexpressing (TG) littermate controls from line-L (white/pink symbols), line-M (white/red symbols), and line-H (white/dark red symbols). Group means are indicated by blue lines. Groups within each individual line were compared by two-way ANOVA to determine significant effects of genotype and sex on data variation and to identify interactions between these variables (P<0.05 denoted in bold red). In the event of significant interaction between genotype and sex, individual groups were compared by Šidak’s multiple comparisons test (NS=P≥0.05, *=P<0.05, **=P<0.01, ***=P<0.001, ****=P<0.0001). **E)** Western blotting of bone marrow total cell lysates from either WT or *Erfe*-overexpressing 10 to 12-month-old, male mice from line-M (TG-M) or line H (TG-H) for pSMAD5, SMAD5, and β actin.

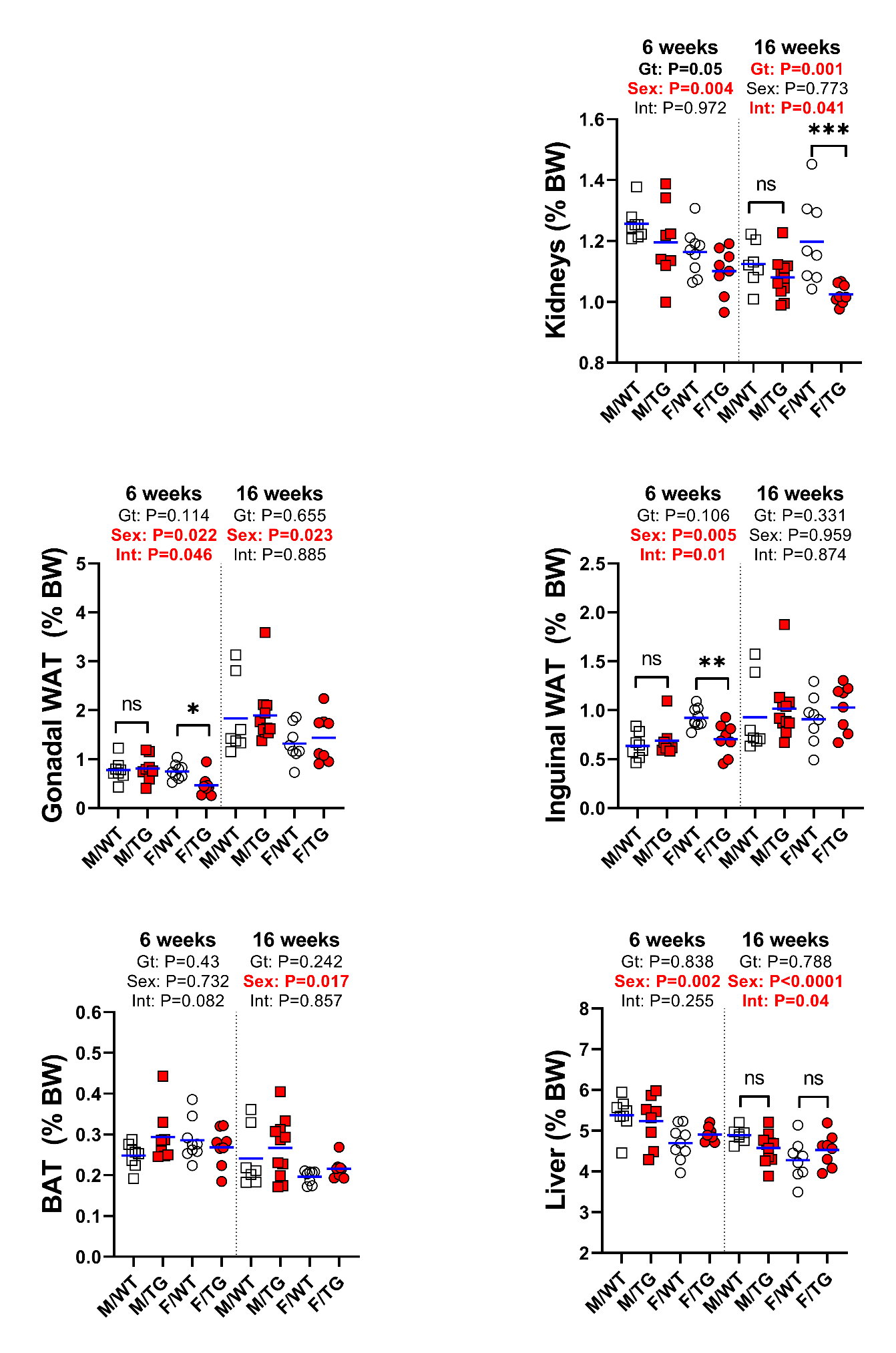

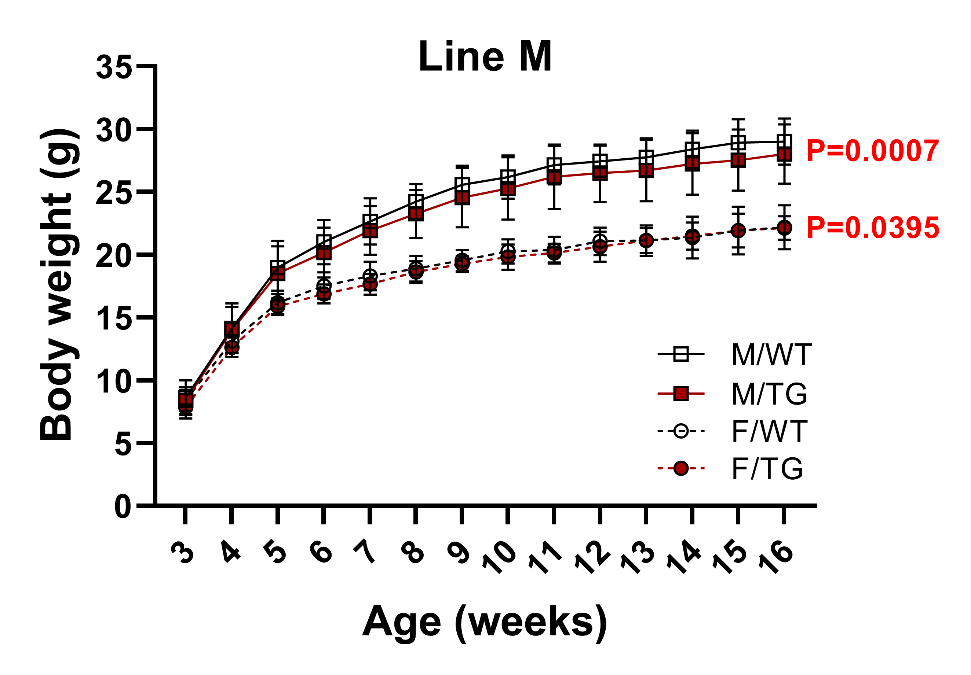
**Supplemental Figure 7**

**B**

**A**

**F**

**E**

**D**

**C**

**Supplemental Figure 7 continued**

**H**

**G**

**I**

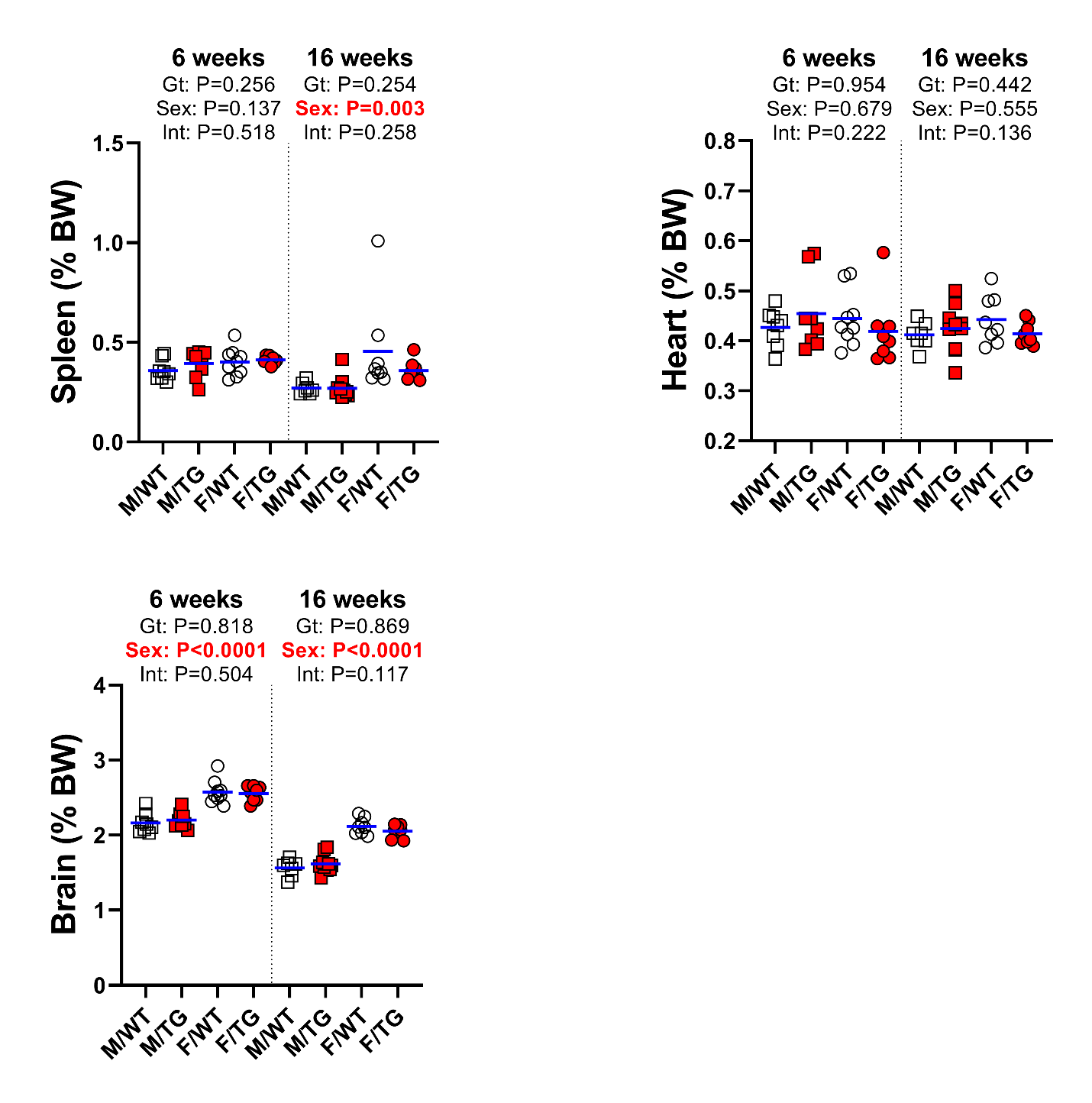

**Supplemental figure 7.** **Growth and tissue mass in line-M mice. A)** Growth curves of line-M mice from ages 3 to 16 weeks of age. Statistical significance of the effect of genotype on data variation between body weights of sex-matched, line-M male (M, square) and female (F, circle) wild-type (WT, white symbols) mice and transgenic (TG, red symbols) littermates determined by 2-way ANOVA are displayed to the right of their respective curves. Statistically significant differences (P<0.05) between sex-matched WT and TG body weights at individual time points, determined by Student’s *t*-test, are indicated by asterisks (*) above or below curves for males and females, respectively. Mass of **B**) kidneys, **C**) gonadal white adipose tissue (WAT) pads, **D**) inguinal white adipose tissue (WAT) fat pads **E**) interscapular brown adipose tissue (BAT), **F**) liver, **G**) spleen, **H**) heart, and **I**) brain at 6 and 16 weeks of age relative to total body mass. Group means are indicated by blue lines. Groups at each age were compared by two-way ANOVA to determine significant effects (P<0.05) of genotype and sex on data variation and to identify interactions between these variables. In the event of significant interaction between genotype and sex, individual groups were compared by Šidak’s multiple comparisons test (NS=P≥0.05, *=P<0.05, **=P<0.01, ***=P<0.001).

**Supplemental Figure 8**

**A**

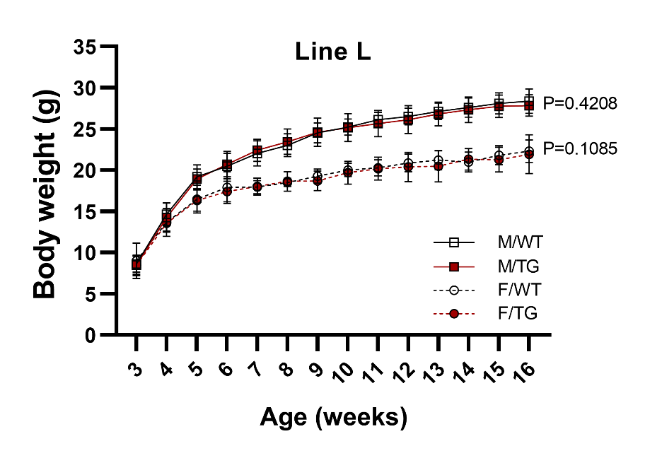

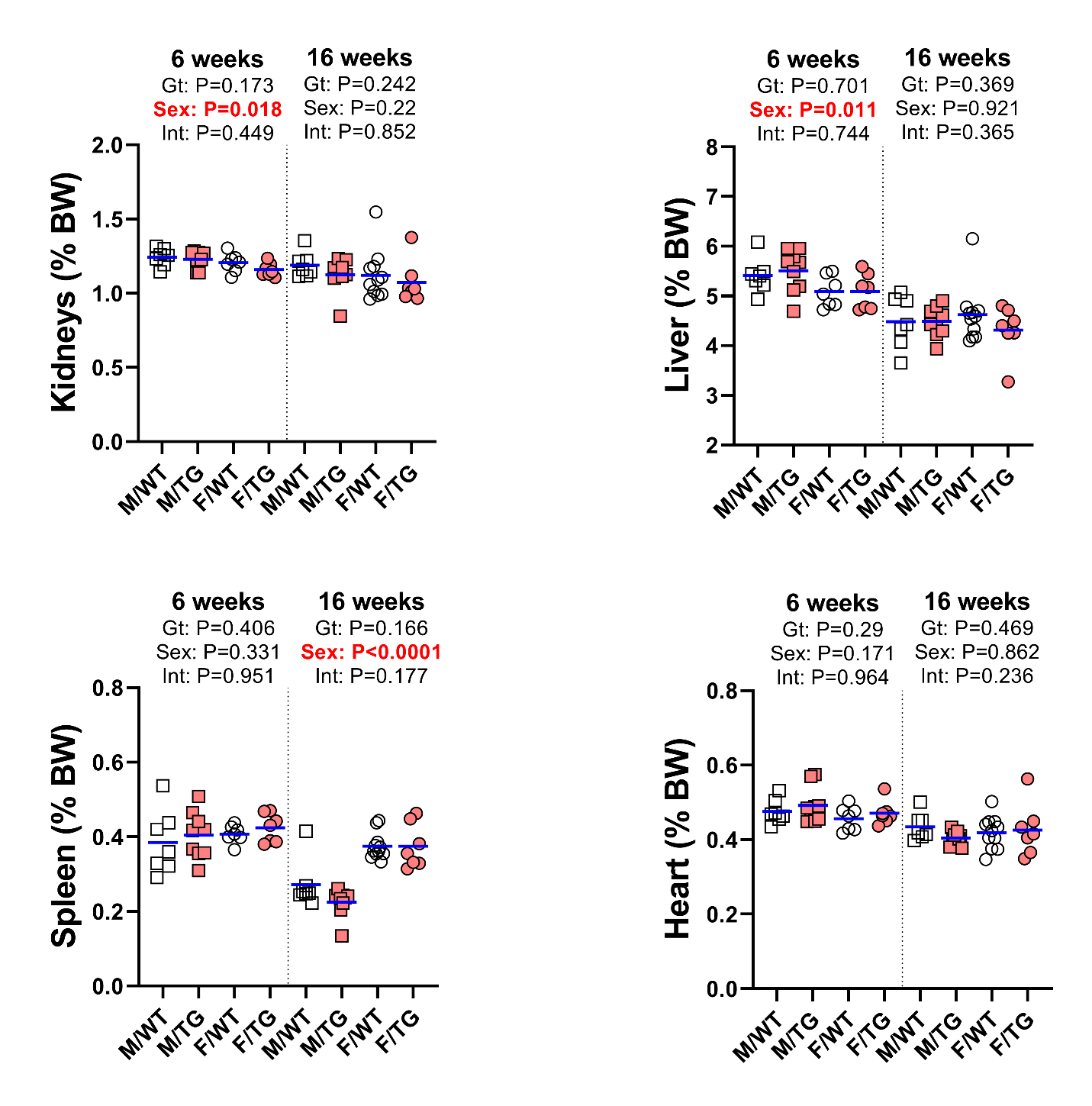

**B**

**C**

**D**

**E**

**Supplemental figure 8. Growth and tissue mass in line-L mice. A)** Growth curves of line-L mice from ages 3 to 16 weeks of age. Statistical significance of the effect of genotype on data variation between body weights of sex-matched, line-M male (M, square) and female (F, circle) wild-type (WT, white symbols) mice and transgenic (TG, pink symbols) littermates determined by 2-way ANOVA are displayed to the right of their respective curves. Statistically significant differences (P<0.05) between sex-matched WT and TG body weights at individual time points, determined by Student’s *t*-test, are indicated by asterisks (*) above or below curves for males and females, respectively. Mass of **B**) kidneys, **C**) liver, **D**) spleen, and **E**) heart at 6 and 16 weeks of age relative to total body mass. Group means are indicated by blue lines. Groups at each age were compared by two-way ANOVA to determine significant effects (P<0.05) of genotype and sex on data variation and to identify interactions between these variables.

**Supplemental Figure 9**

**B**

**A**

**C**

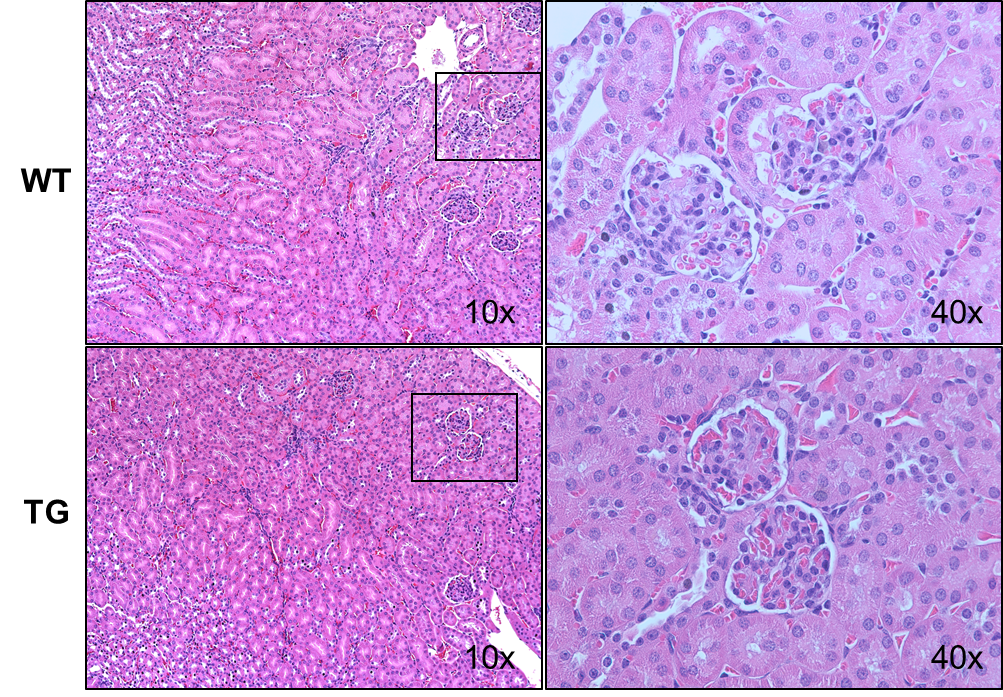

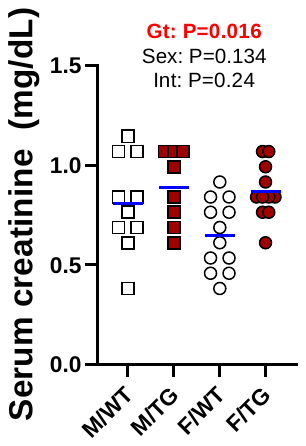

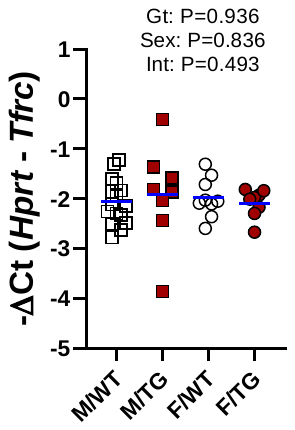

**Supplemental figure 9. Characterization of additional kidney-associated parameters in line-H mice. A)** Serum creatinine levels in male (M, square) and female (F, circle) wild-type (WT) mice and *Erfe*-overexpressing (TG) littermates from line-H (white/dark red) at 16 weeks of age. **B)** Representative hematoxylin and Eosin staining of kidney sections from 6-week-old, male, WT and TG mice from line-H at 10x and 40x magnification. Kidney regions selected for 40x images are indicated by black outlines on corresponding 10x images. **C)** Relative mRNA expression of *Tfrc* in kidneys from 6-week-old, line-H mice. Group means are indicated by blue lines and groups within each individual line and age group were compared by two-way ANOVA to determine significant effects of genotype and sex on data variation and to identify interactions between these variables (P<0.05 denoted in bold red) and to identify interactions between these variables.

**Supplemental Figure 10**

**A**

**B**

**D**

**C**

**E**

**F**

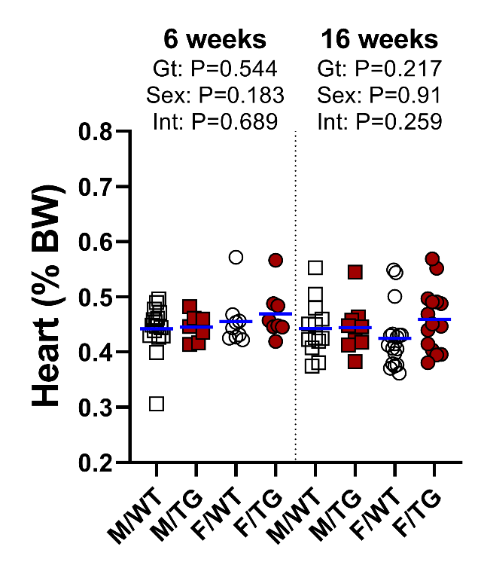

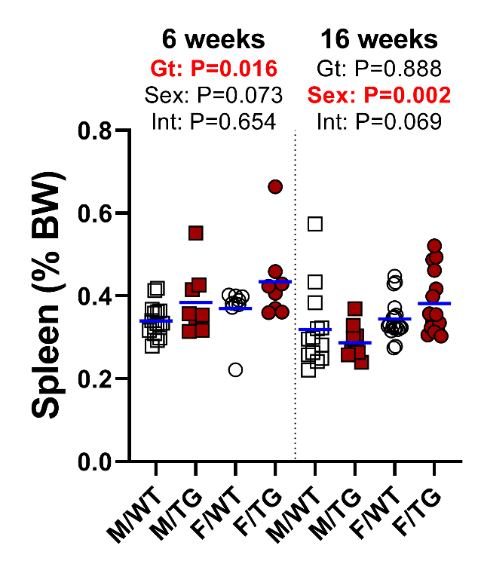

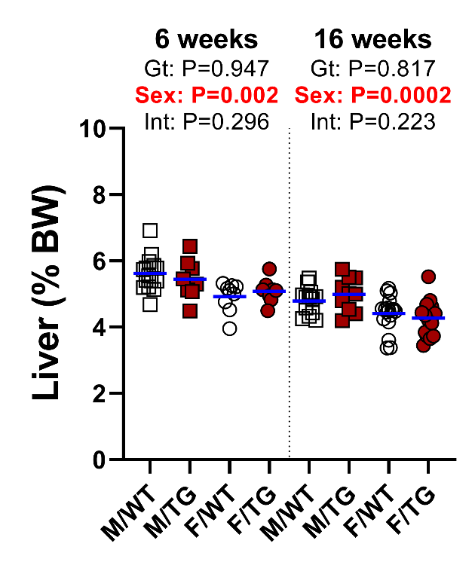

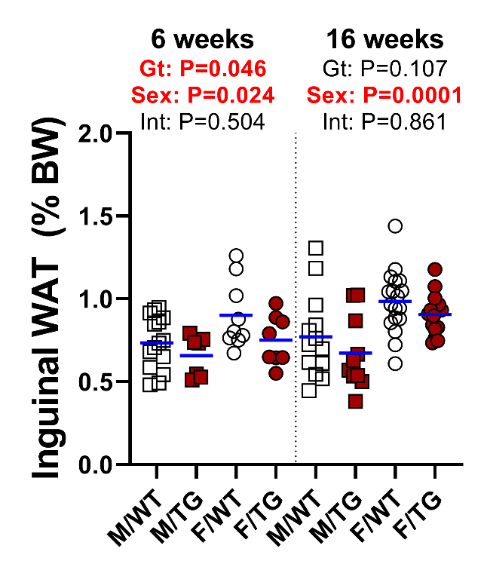

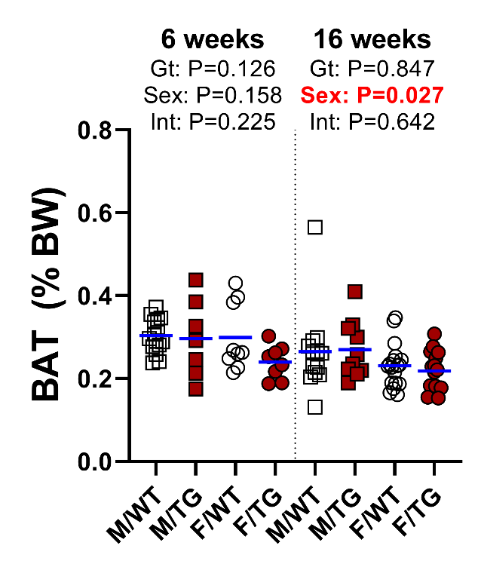

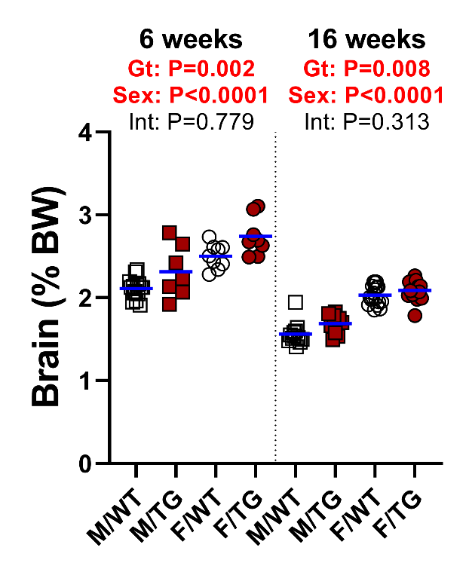

**Supplemental figure 10. Mass of various tissues from line-H mice.** Mass of **A**) brain, **B**) inguinal white adipose tissue (WAT) fat pads, **C**) interscapular brown adipose tissue (BAT), **D**) liver, **E**) spleen, and **F**) heart at 6 and 16 weeks of age relative to total body mass in line-H male (M, square) and female (F, circle) wild-type (WT, white symbols) mice and transgenic (TG, dark red symbols) littermates. Group means are indicated by blue lines. Groups at each age were compared by two-way ANOVA to determine significant effects (P<0.05) of genotype and sex on data variation and to identify interactions between these variables.

**Supplemental Figure 11**

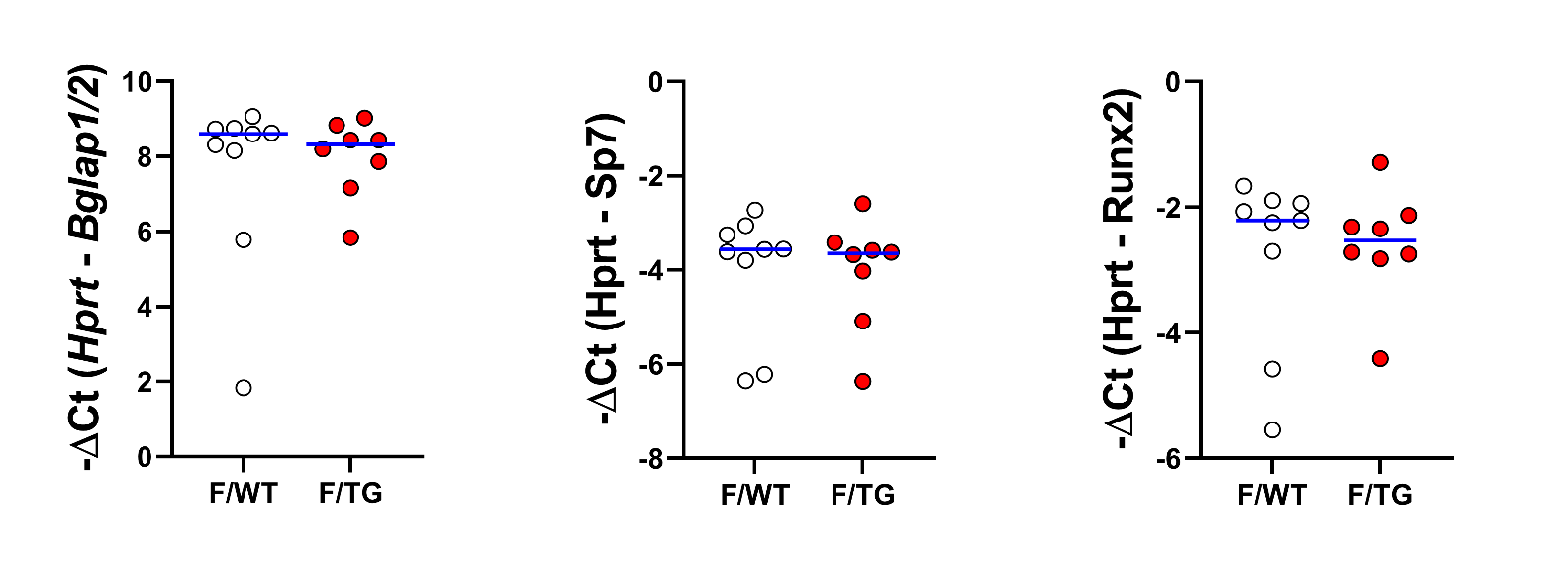

**A**

**B**

**C**

**Supplemental figure 11. Effect of *Erfe* overexpression on the expression of bone-associated genes.** Relative mRNA expression of **A)** *Bglap1/2*, **B)** *Sp7*, **C)** *Runx2* in tibia bone at 6 weeks of age in female (F) wild-type (WT) mice and *Erfe*-overexpressing (TG) littermate controls from line-M (white/red symbols). Group means are indicated by blue lines. Differences in group means between F/WT and F/TG mice were analyzed for statistical significance by Student’s *t*-test (*=P<0.05).

**Supplemental Figure 12**

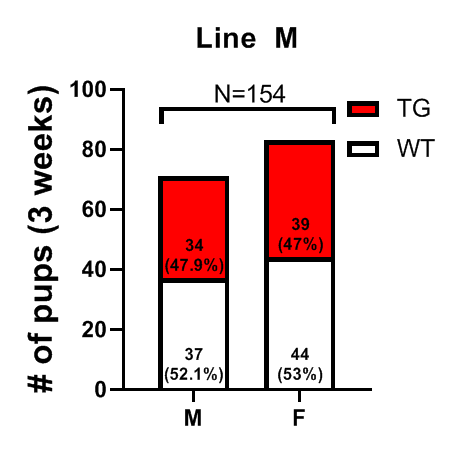

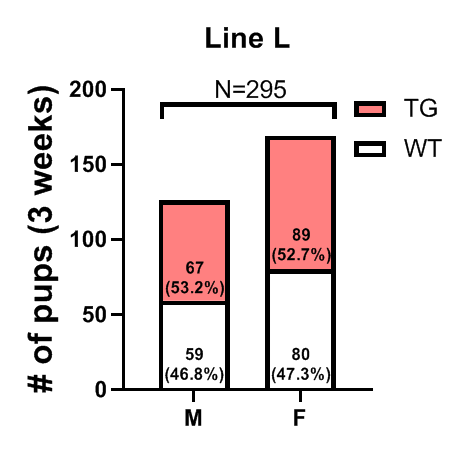

**A**

**B**

**Supplemental figure 12. Normal postnatal survival of line-L and line-M transgenic mice.** Number and within-sex percentage of wild-type (WT, white) and *Erfe*-overexpressing (TG) mice, grouped by offspring sex (male=M, female=F), from **A**) line-L (pink) and **B**) line-M (red) mice surviving at 3 weeks of age. Differences from expected proportions were not statistically significant by binomial testing at (P<0.05).

**Supplemental Figure 13**

**A**

**B**

**C**

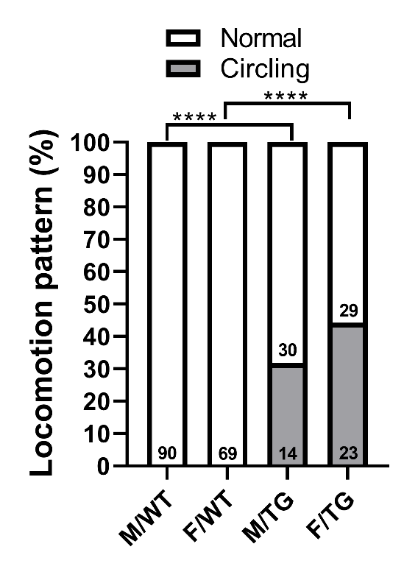

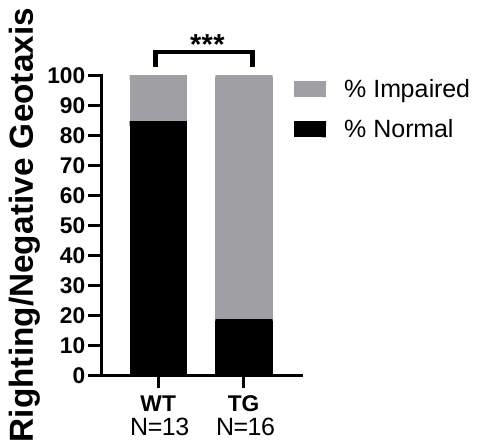

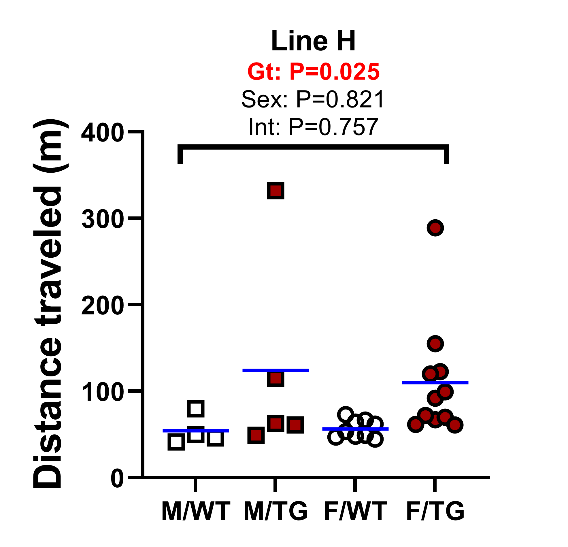

**Supplemental figure 13.** **Repetitive circling behavior and impaired motor/vestibular function in line-H transgenic mice. A**) Incidence of overt circling behavior in line-H mice. Numerical values on bars indicate animal number per pattern of movement in male (M) and female (F) wild-type (WT) mice or *Erfe*-overexpressing (TG) littermates. Statistically significant differences in locomotion pattern between groups of the same sex were determined by Fisher’s exact test (****=P<0.0001). **B)** Total distance traveled during open-field testing by line-H mice. Group means are indicated by blue lines. Groups were compared by two-way ANOVA to determine significant effects of genotype and sex on data variation and to identify interactions between these variables (P<0.05 denoted in bold red). **C**) Incidence of impaired outcomes during righting reflex or negative geotaxis testing in mixed-sex, line-H WT and TG mice. A mouse was designated as “impaired” if its combined score from the righting reflex and negative geotaxis testing was >0. Mice categorized as normal had a combined score of 0 on these tests. Statistical significance of the difference in incidence of impairment between groups was determined by Fisher’s exact test (***P<0.001).
